# A semantics, energy-based approach to automate biomodel composition

**DOI:** 10.1101/2021.11.12.468343

**Authors:** Niloofar Shahidi, Michael Pan, Kenneth Tran, Edmund J. Crampin, David P. Nickerson

**Affiliations:** Auckland Bioengineering Institute, The University of Auckland, Auckland, New Zealand; Systems Biology Laboratory, School of Mathematics and Statistics, and Department of Biomedical Engineering, University of Melbourne, Melbourne, Victoria, Australia; ARC Centre of Excellence in Convergent Bio-Nano Science and Technology, Faculty of Engineering and Information Technology, University of Melbourne, Melbourne, Victoria, Australia; School of Mathematics and Statistics, Faculty of Science, University of Melbourne, Victoria, Australia; School of Medicine, University of Melbourne, Victoria, Australia

## Abstract

Hierarchical modelling is essential to achieving complex, large-scale models. However, not all modelling schemes support hierarchical composition, and correctly mapping points of connection between models requires comprehensive knowledge of each model’s components and assumptions. To address these challenges in integrating biosimulation models, we propose an approach to automatically and confidently compose biosimulation models. The approach uses bond graphs to combine aspects of physical and thermodynamics-based modelling with biological semantics. We improved on existing approaches by using semantic annotations to automate the recognition of common components. The approach is illustrated by coupling a model of the Ras-MAPK cascade to a model of the upstream activation of EGFR. Through this methodology, we aim to assist researchers and modellers in readily having access to more comprehensive biological systems models.

**Author summary:** Detailed, multi-scale computational models bridging from biomolecular processes to entire organs and bodies have the potential to revolutionise medicine by enabling personalised treatments. One of the key challenges to achieving these models is connecting together the vast number of isolated biosimulation models into a coherent whole. Using recent advances in both modelling techniques and biological standards in the scientific community, we developed an approach to integrate and compose models in a physics-based environment. This provides significant advantages, including the automation of model composition and post-model-composition adjustments. We anticipate that our approach will enable the faster development of realistic and accurate models to understand complex biological systems.

## 1 Introduction

Modelling complex biological systems such as cells, organs, and organisms allows researchers and physicians to integrate and study different aspects of a biological entity, reveal the limits and shortcomings of our knowledge, and obtain new insights into disease treatment [1]. Motivated by such aims, hierarchical modelling is an approach that assists researchers in constructing system-level models, which are continuously expanding in detail, scope, and size [2].

Hierarchical models are composed of pre-existing smaller models, referred to as modules. Each module can operate and be examined independently, thus reducing model composition errors and facilitating large-scale model generation due to pre-existing models. To accelerate hierarchical model composition, one can capitalise on a myriad of the existing modules created by others. A requirement for this is that the modules be both accessible and reusable [2]. Over the past decade biosimulation models have become increasingly accessible on public repositories such as the Physiome Model Repository (PMR) [3] and BioModels [4], which store models in XML-based format such as CellML [5] and the Systems Biology Markup Language (SBML) [6, 7].

The main challenges in hierarchical model composition include: (a) incompatible code languages, (b) different modelling frameworks, (c) post-composition adjustments, and (d) physically implausible resultant models. Each integrating system or platform addresses some of these challenges. A majority of model integration platforms, such as the SBML Hierarchical Model Composition package [8], require compatibility between the languages and modelling frameworks in the modules. In contrast, the resultant model still needs further post-merging code-wise adjustments to be executable, yet it might not represent a physically feasible model. Physical feasibility refers to models following the laws of thermodynamics and physics even if the model itself is incorrect [9, 10]. One solution to these issues is using a hierarchical modelling approach (to help with the post-composition adjustments) and an energy-based modelling framework (to guarantee a physically plausible composed model). The bond graph approach addresses these issues.

Bond graphs provide a domain-independent hierarchical framework that generates models based on the laws of physics and thermodynamics. Initially introduced by Paynter [11], bond graphs were primarily intended for engineering applications. The application of bond graphs was extended to the chemical domain by Oster et al. [12, 13] and subsequently by Cellier [14]. Gawthrop and Crampin have recently developed the bond graphs framework to model and analyse biochemical and electrochemical systems [10, 15].

An automated model composition approach significantly assists researchers in creating large-scale models from existing modules [16]. Shahidi et al. [17] introduced a general hierarchical model composition method by encoding bond graph modules in CellML and constructing a complex model using the semantics-based SemGen merger tool [18]. Although this method facilitated the integration of annotated bond graph models, bottlenecks might arise when a modification in the CellML bond graph modules is needed (modellers must know the bond graph conservation laws). Moreover, it required adding auxiliary variables as ports to each module and connecting them manually using the semi-automated SemGen merger tool. While annotations are readily incorporated into bond graphs, using annotations in model composition has not been conducted in this context.

Here, as an extension to our previous work, we have incorporated annotations to bond graphs in a platform in which a composed model can be automatically constructed from annotated CellML files treated as modules. Because the CellML files do not contain the bond graph structure, a separate bond graph library in Python (BondGraphTools [19]) automatically deals with any required changes in the conservation laws. The annotated data from the CellML modules are then extracted and assigned to their equivalent bond graph components. Thereafter, any common entities among the modules are identified and merged to render a composed model. We demonstrate this by an example where a bond graph model is constructed from its constitutive modules, *i.e.*, the Epidermal Growth Factor Receptor (EGFR) signalling pathway and the Mitogen-Activated Protein Kinase (MAPK) cascade are merged to construct a model of the entire EGFR-Ras-MAPK signalling pathway. This type of model integration provides a reliable and consistent framework that first conserves energy due to the energy-based bond graphs implementation; second, prevents modellers from making physically infeasible models; third, does not require any post-merging modifications due to the hierarchical feature of bond graphs; and fourth, automates the model composition and merging using the modules’ rich semantics.

In this paper, we introduce our automated model composition approach in Section 2.1. Its prerequisites along with the generic method description are reviewed in Sections 2.1.1 & 2.1.2. We review the use of bond graphs in modelling biochemical reactions (Section 2.2) and describe how the bong graph modules of the EGFR-Ras-MAPK signalling pathway are created based on the existing work by Kholodenko et al. [20] and Pan et al. [21] (Section 2.3). As an example of our method application, we utilise it to create the EGFR-Ras-MAPK signalling pathway in Section 2.4. Next, we describe how we verified the simulation results of our composed model in Section 2.5. In Section 3 we demonstrate the simulation results both for the constitutive modules and our composed bond graph model, and in Section 4 we discuss and analyse the behaviour of bond graph modules and then verify the behaviour of our composed model. Possible improvements and shortcomings are also discussed in this section. The main features of our method and the future developments are summarised in Section 5.

## 2 Materials and methods

In this section, we discuss our energy-conserving semantics-based model composition approach: the prerequisites and the generic application. We give a brief introduction to bond graph modelling of biochemical networks consisting of multiple reactions. Later, we demonstrate how a mathematical model of a biochemical network can be converted into bond graphs. We utilised this approach to create a bond graph model of the EGFR pathway. We will show that by having the bond graph model of any physical or chemical system, our method can merge the similarly annotated entities within the models by automatically rewiring the connections between components and modules. The whole composed model in bond graphs will then be ready for simulation or connection to other modules. We demonstrate this by applying our method to automatically compose and generate a EGFR-Ras-MAPK signalling model.

### 2.1 Automated model composition pipeline

Our method to expand and integrate biosimulation models provides the foundation for further developments in an open-source environment based on energy-based modules and automation to minimise manual input. In this endeavour, we have provided some exemplar symbolic bond graph models to which the annotated parameters from the CellML modules would accordingly link. Symbolic modules are predefined models in which the parameters do not have any values and allow us to determine the parameters’ values later where each module gets its specific parameters from the source CellML files. The suitable bond graph model is then automatically selected from the list by identifying specific annotated components in the CellML modules. Merging points are automatically recognised and merged, resulting in a physically consistent model. Due to the hierarchical feature of bond graphs, the needed adjustments during the composition will be systematic, leading to the automation of modifications (adding/deleting bonds between the components).

To apply our method to CellML models, some preparations are required in advance, *i.e.*, installing the bond graph Python library as well as downloading the required ontologies. Once prepared, the user can commence the model composition in a Python environment like Jupyter Notebook.

#### 2.1.1 The prerequisites

• BondGraphTools

The task of automated model composition requires bond graph software that readily supports automation. For this purpose, we have selected BondGraphTools

– an open-source Python library for bond graph modelling – created and developed by Cudmore et al. [19], accessible from https://github.com/BondGraphTools/BondGraphTools. BondGraphTools supports modularisation and automation in model building.

• Ontologies

Depending on the area of biomedical science in which the researchers annotate their models, one or various reference ontologies might be used. Our approach identifies and interprets the annotations by comparing them with the codes and labels in ontologies saved as *csv* files. We suggest downloading CHEBI, FMA, OPB, and GO ontologies from the following links:

**–** CHEBI: https://bioportal.bioontology.org/ontologies/CHEBI
**–** FMA: https://bioportal.bioontology.org/ontologies/FMA
**–** OPB: https://bioportal.bioontology.org/ontologies/OPB
**–** GO: https://bioportal.bioontology.org/ontologies/GO

We used the OPB and GO ontologies for the particular case study in this paper.

#### 2.1.2 The generic approach

To reuse and compose models deposited on online repositories such as PMR, we need a tool to first convert a non-bond-graph model into an equivalent bond graph one; second, automatically assign the parameters in the models to the bond graph components; third, identify the same entities in the models as the merging points and make the necessary changes to join the models without any loss of information.

To improve our model composition method toward automation and reuse the models in various formats, we employed the idea of having symbolic bond graph templates and connectivity matrices for some exemplar systems (EGF signalling and the Ras-MAPK cascade). A connectivity matrix is a binary square matrix that defines connections between the elements of a system. The number of rows and columns each equals the number of elements in the network in total [22]. Instead of using the embedded syntax in BondGraphTools to append/delete the elements, we employed the concept of a connectivity matrix. Binary representation of models clearly shows the connections, facilitates computational measurements, and gives the minimal required details to define a network which can be exported to other tools and software for further analysis [22]. Modifying a network is easily performed by inserting 0 or 1 in the matrix or deleting its corresponding row and column. The connectivity matrix (*CM* ) for a system with *m* elements can be described as *CM ∈ {*0, 1*}_m×m_*, where:

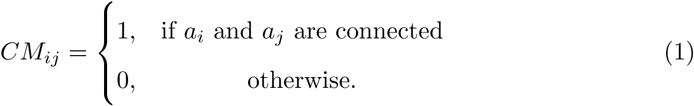

Figure 1 shows the connectivity matrix for a sample network. Notice that the connectivity matrix is symmetric (*CM_ij_*= *CM_ji_*), representing the bidirectional flow of energy between the bond graph components.

**Fig 1.**
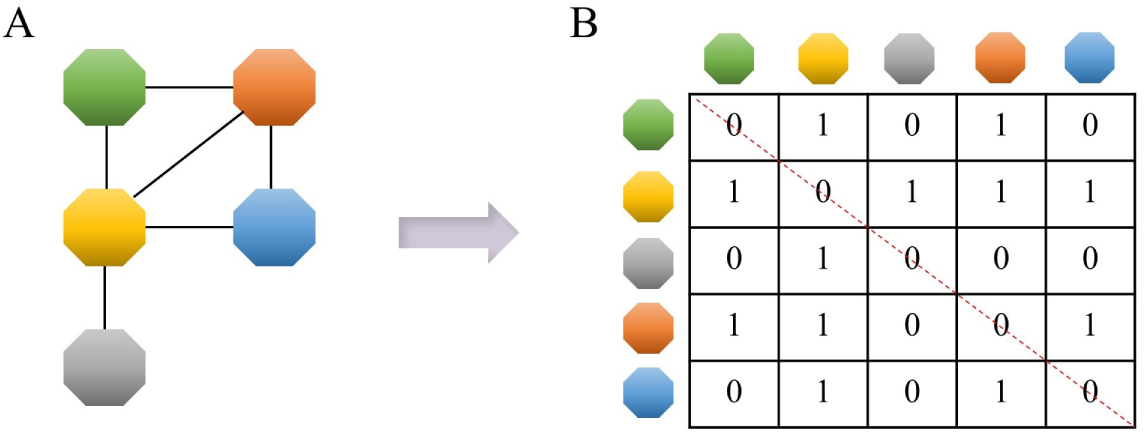
An example network with its connectivity matrix. (A) The network topology of the connections between the elements of a system; (B) The connectivity matrix of the system. 0 means *no connection* and 1 means *connection*. The dashed red line shows the diagonal of the matrix to demonstrate its symmetry.

To identify the merging points between the modules we used a ‘white box’ approach. In this approach all or a group of the elements in the modules can be selected as merging points. In a ‘black box’ composition approach, the elements of the modules are not accessible and only the input/output variables can be used for coupling modules [17, 23]. In coupling biological models almost all the entities can be regarded as merging ports, hence, we found the ‘white box’ configuration more compatible with our model composition method. To do this, we need the variables and parameters of the models to be annotated. To summarise, we started our automated model composition method by preparing the following (Figure 2):

- Bond graph symbolic templates of the models;
- Connectivity matrix of each bond graph template;
- Ontologies required for matching and interpreting the annotations.

**Fig 2.**
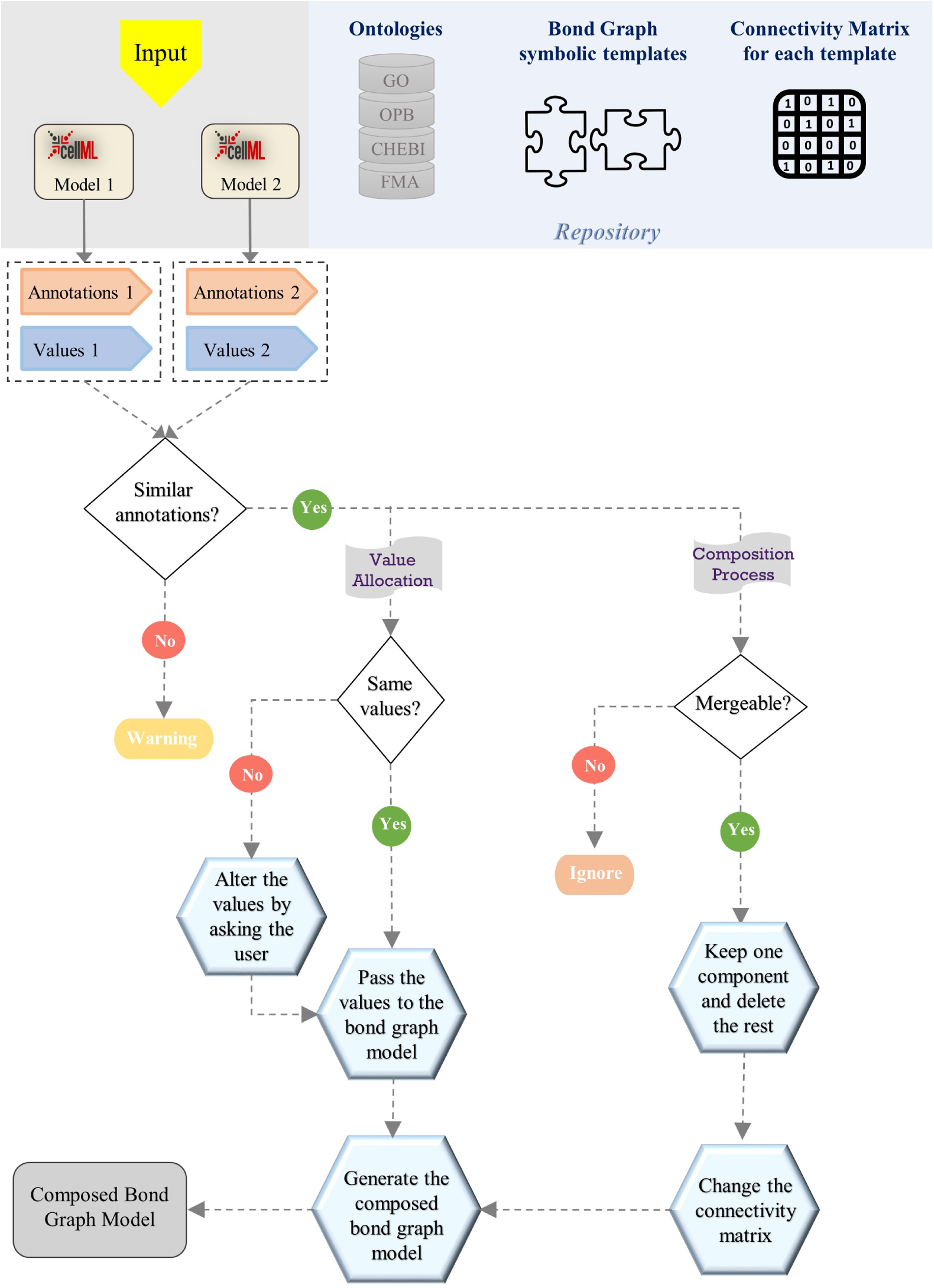
The generic flowchart of our automated model composition approach. The Repository shows the prerequisites for model composition. The Input section shows two arbitrary CellML models to be merged using our approach.

In a given biological/physiological/physical context, our framework can detect the type of bond graph template that matches the annotated model. This is done by searching for specific groups of biological entities/processes within the annotated CellML files. If a certain group of entities is found in a file, then it will be linked to its corresponding bond graph symbolic template. Thereafter, by finding similar annotations in the models, the merging points will be selected. Based on this, the required changes in the bond graph components (deleting the duplicates) and the connectivity matrices (deleting or inserting rows and columns) will be made. Ultimately, the final model will be produced based on the connection/non-connection relationships between all the components.

We have deposited the required ontologies, the bond graph symbolic template models, and their connectivity matrices in our repository. Any number of CellML models containing the annotated parameters of a system can be used in our framework as inputs (here, we list two models). First, the annotations and values of the CellML models are extracted, and if no similar annotations are detected between the models, it gives a warning. For the similar annotations, two paths are taken:

1. Value allocation: If the same components do not have the same values, it asks the user to either select one of the values or insert a new one.
2. Composition process: If the components with similar annotations are mergeable, it keeps one of them and discards the rest; otherwise, it ignores them [24]. For example, biochemical species are considered mergeable since they can simultaneously participate in multiple reactions but a parameter like temperature cannot be merged as it can not become a port for external connections. Based on the deleted duplicate components, the connectivity matrices are combined, allowing the models to be merged (details in Section 2.4).

### 2.2 Bond graph modelling of biochemical reactions

This section summarises the bond graph basic principles and delineates how biochemical reactions are represented in bond graphs.

Two physical co-variables form the energy-based foundation of bond graphs: *effort* (*e*) and *flow* (*f*). Power is the product of effort and flow (*p* = *e.f* ) and energy is the power over time: *E* = ∫*p dt*. Effort and flow are general terms that represent voltage and current in the electrical domain, force and velocity in the mechanical domain, chemical potential and molar flux in the chemical domain, respectively. Bond graphs represent complex systems as graphical representations which consist of components, bonds, and junctions. Components represent physical elements and are defined as general configurations of electrical, mechanical, or chemical elements. For instance, *C* components in bond graphs are charge storage components i.e. capacitors in electrical circuits, springs in mechanical systems, or chemical species in chemical reactions. A common effort between the components is shown by a ‘0’ junction, while a ‘1’ junction shows a common flow, and the energy is conserved and travels between components bidirectionally through bonds (shown by harpoons). Readers can find a more detailed description of the bond graph theory in [25–27].

To facilitate reusability, biochemical models must obey the laws of physics and thermodynamics [28]. In conventional modelling approaches, modellers often ignore the energy transfer; thus, the reactions may proceed against chemical potential gradients and lead to physically implausible models [10]. Since bond graphs are based on energy conservation and thermodynamic laws, fluxes are always in the direction of decreasing potential. Biochemical bond graph models contain components for the species (*C_e_*), stoichiometry (*TF* : *N* ), and reaction (*Re*). To highlight the notion of the bond graph junctions for sharing a *common molar flux* or a *common chemical potential*, we indicate them by ‘1 : *v*’ and ‘0 : *u*’, respectively. Here,

- The chemical potential is *u* (J mol*^−^*^1^), stored within the biochemical species, and the molar flux is *v* (mol s*^−^*^1^), driven by the reactions;
- The biochemical species are defined using the component *C_e_*, given by the constitutive relation *u_q_* = *RT* ln *K_q_.q* (**Boltzmann’s** formula), where *R* (J mol*^−^*^1^ K*^−^*^1^) is the ideal gas constant, *T* (K) is the temperature, *q* (mol m*^−^*^3^) is the molar concentration of the species, and *K_q_* (mol*^−^*^1^) is the thermodynamic constant of the species [29];
- Species with fixed concentrations are considered as sources of potential which are called chemostats (*C_S_* ) in bond graph terminology;
- A reaction represents a dissipative process, which in the case of mass-action kinetics is defined by an *Re* component with the constitutive relation *v* = *κ*(*e^ur/^*^(^*^RT^*^)^*− e^up/^*^(^*^RT^*^)^) (**Marcelin–de Donder** equation), where *κ* is the reaction rate constant and *u_r_* and *u_p_* are total chemical potentials of the reactants and products, respectively;
- Stoichiometries are represented by transformer *TF* : *N* , in which the transformer ratio (*N* ) corresponds to stoichiometry.

For further discussion of bond graph modelling of biomolecular and chemical systems, the reader is referred to the works by Gawthrop & Crampin [21, 30].

As an example, a reaction with two reactants and two products is demonstrated in Fig 3 along with its equivalent bond graph representation.

**Fig 3.**
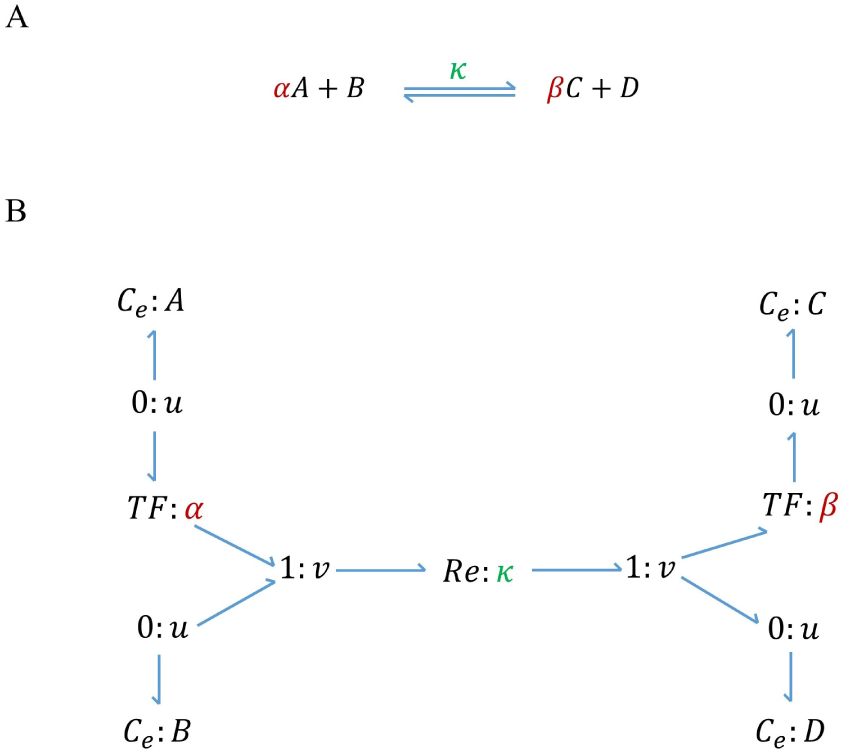
A chemical reaction and its bond graph equivalent. A chemical reaction with two reactants and two products. (A) Schematic of a chemical reaction where *κ* is the reaction rate constant, *A* & *B* are the reactants, *C* & *D* are the products, and *α* & *β* are stoichiometries; (B) Bond graph equivalent of the reaction where *C_e_* components correspond to the species, *Re* corresponds to the reaction, and *TF* components represent the stoichiometries. Since the consumption/production rate of all the contributing species in a reaction is equal to the reaction flow rate, they share a common flow with the *Re* component through a ‘1 : *v*’ junction.

In Fig 3.B, the reaction flow rate for the *Re* component is given by:

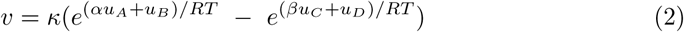

or if we substitute the chemical potentials with the Boltzmann’s formula:

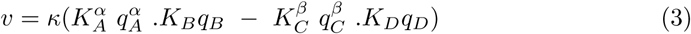

which can be generally described by mass action kinetics:

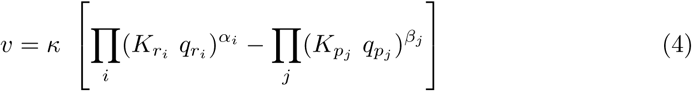

where *K_ri_*and *K_pj_*are the thermodynamic constants, *q_ri_*and *q_pj_*are the concentrations, and *α_i_* and *β_j_* are the stoichiometries of reactants and products, respectively.

In the next section we illustrate how the bond graph approach toward biochemical reactions is utilized to create models of two exemplar biochemical pathways.

### 2.3 Modules for EGFR-Ras-MAPK signalling: Bond graph models of the pathways

The EGFR-Ras-MAPK is a signalling pathway that transduces signals from the extracellular environment to the cell nucleus [31]. It participates in multiple biological functions in mammalian cells, including growth and differentiation, cell migration, and wound healing [20, 31, 32] and consists of two major parts: the EGFR pathway and MAPK cascade. Ras protein activation signals stimulate the Ras-MAPK cascade [20], which is located downstream at the EGFR pathway. As such, we consider Ras protein to be the mutual species in both pathways. The bond graph model of the EGFR pathway is created based on the model by Kholodenko et al. [20] (CellML model available from: *Kholodenko 1999* ) and the bond graph model of MAPK cascade is taken from the work by Pan et al. [21]. In this paper, the bond graph representation of the reference MAPK cascade was available. Here, we detail how bond graph models of these systems were constructed.

#### 2.3.1 EGFR pathway module

The schematic of the EGFR pathway model developed by Kholodenko et al. [20] is shown in Fig 4. The signal transmission starts with the Epidermal Growth Factor (EGF) binding to the Epidermal Growth Factor receptor (EGFR). It continues via several subpathways that target the SOS (Son of Sevenless) protein. The formation of SOS complexes (RShGS and RGS) activates the Ras protein, which initialises phosphorylation in the MAPK cascade.

**Fig 4.**
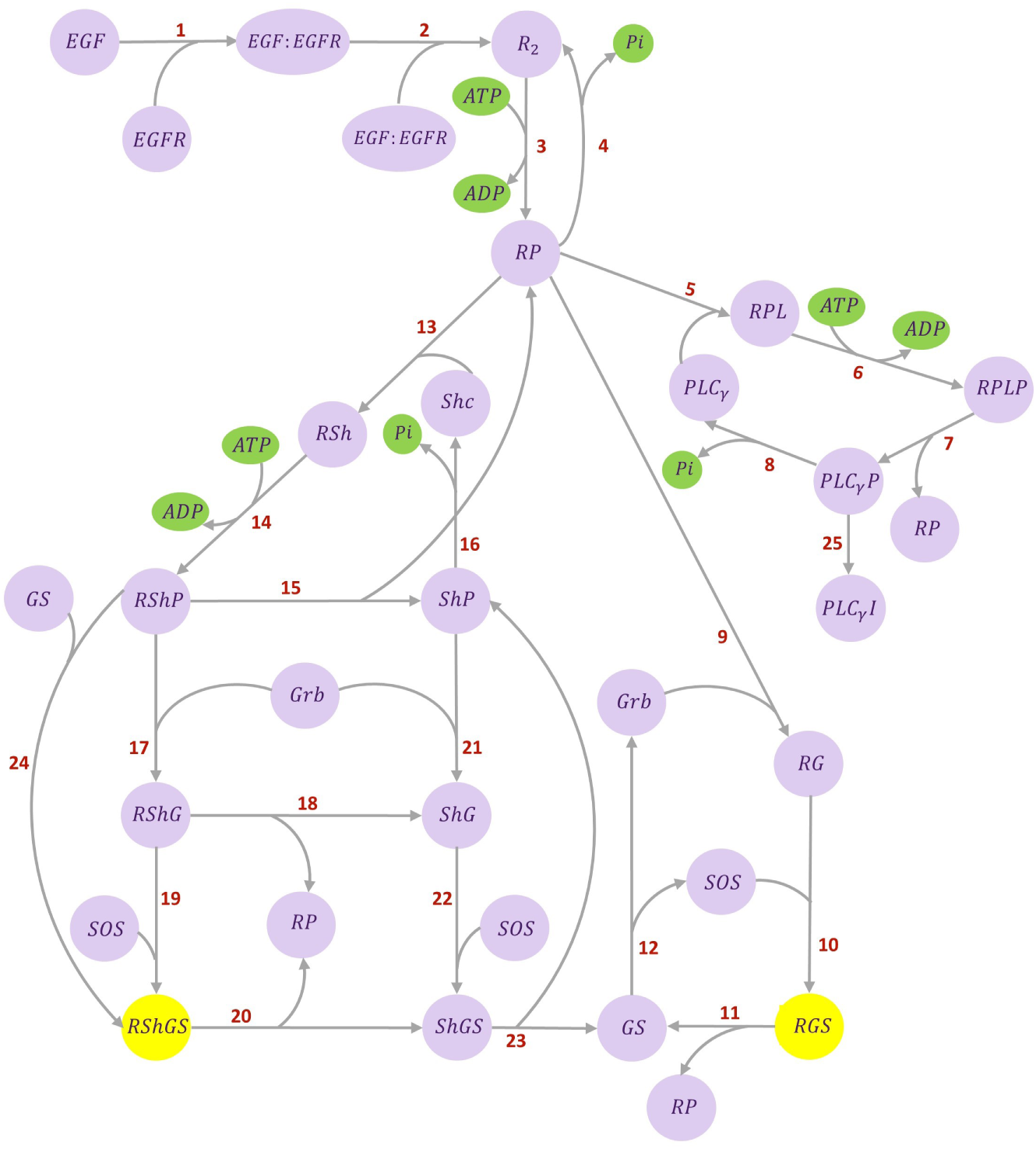
Kinetic structure of EGFR pathway. The ATP hydrolysis species are shown in green (involved in phosphorylation and dephosphorylation reactions). The complexes in yellow activate the Ras protein. The reactions are numbered as the equations in the CellML source code. The network adapted from [20].

The bond graph equivalent network of the EGFR pathway is illustrated in Fig 5.

**Fig 5.**
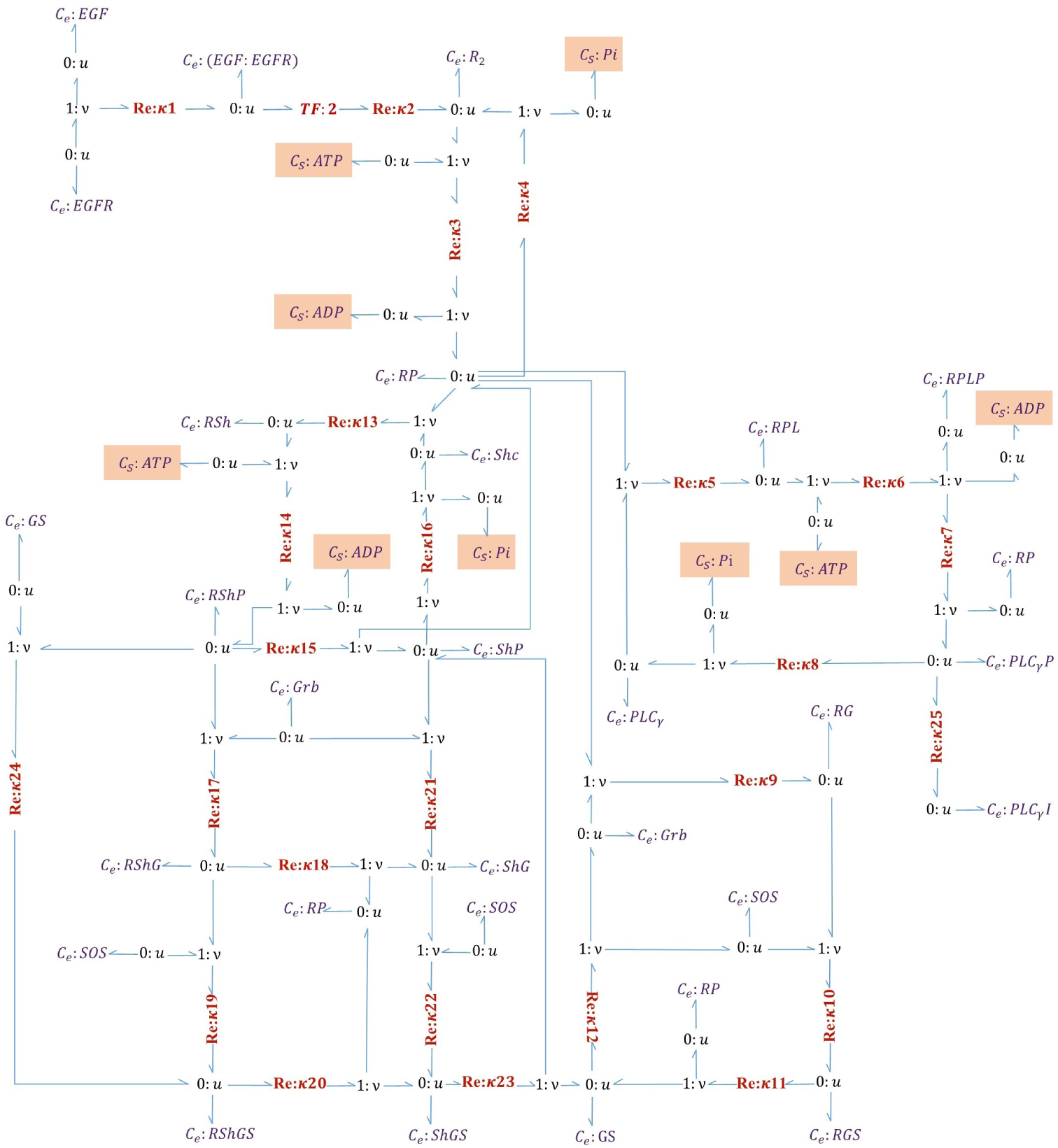
Bond graph representation of EGFR pathway. *Re* components are numbered according to the steps in [20]. Each *C_e_* or *C_S_* component is connected to a ‘0 : *u*’ junction. Where a species participates in more than one reaction, new bonds are applied to its corresponding ‘0 : *u*’ junction to share a common chemical potential (See R-PL where it is produced in reaction 5 and consumed in reaction 6). The chemostats in orange boxes are added to the reconstructed bond graph version.

The reactions in the EGFR model by Kholodenko et al. are either reversible or irreversible. The reversible reactions are described using the kinetic scheme as:

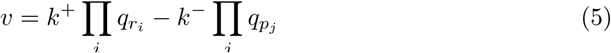

where *k*^+^ and *k^−^* are the forward and reverse kinetic rate constants and Π*_i_ q_r_* and Π*_j_ q_p_* are the concentrations of reactants and products, respectively. The kinetic model and parameters are given in [20] (Table I & Table II). The irreversible reactions (steps 4, 8, 16) are described using **Michaelis–Menten** kinetics as:

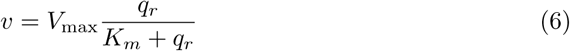

where *V*_max_ (mol*/*s) is the maximum reaction rate achieved by the system, *K_m_* (mol) is the Michaelis constant referring to the reactant concentration at half of the *V*_max_, and *q_r_* (mol) is the reactant concentration. Irreversible reactions are thermodynamically impossible in a bond graph model; we deal with this issue later.

The required energy for the reactions is supplied by Adenosine triphosphate (ATP) hydrolysis, producing Adenosine diphosphate (ADP) during phosphorylation and phosphate (Pi) during dephosphorylation. The reversible phosphorylation reactions (steps 3, 6, 14) follow the kinetic formulation (Eq 5) and the irreversible dephosphorylation reactions (steps 4, 8, 16) follow the Michaelis–Menten kinetics (Eq 6). Kholodenko et al. have not explicitly included ATP, ADP, and Pi in their model, which contravenes mass and energy conservation. Therefore, we considered these species in our bond graph model. ATP, ADP, and Pi are assumed to be chemostats.

To convert the kinetic parameters (*k*^+^ and *k^−^* in Eq 5) to those required by bond graphs (*κ*, *K_r_*, and *K_p_* in Eq 4), we first removed the thermodynamically infeasible irreversible reactions from the network (for their different parameter definitions) and then applied the method described in [28]. In brief, by taking logarithms on the constraints of each reaction (*k*^+^ = *κΠ_i_ K_r_* and *k^−^* = *κ Π_j_ K_p_* ), the relationship between the kinetic and bond graph parameters can be expressed as a linear matrix [28]. Due to accounting for the ATP, ADP, and Pi components in our bond graph model, we included them in the constraints of their corresponding reactions.

As our selected pathways are in the cytosolic compartment, they use the same sources of potential (ATP, ADP, and Pi); thus, we used the same chemical potentials for these chemostats as we used in the MAPK cascade module. We obtained the thermodynamic constants in the previous phase (except the ones in the irreversible reactions), in which we converted the kinetic parameters into bond graph parameters. We approximated the irreversible reactions with kinetic quantities (Eq 5) which led to a negligible reverse molar flux. We applied curve fitting to estimate the reaction rate constants for the irreversible steps (*κ*_4_, *κ*_8_, & *κ*_16_). We obtained the time-dependent behaviour of the contributing species in steps 4, 8, and 16 (required for curve fitting) from the reference CellML model for the EGFR pathway. As we will discuss further in Section 3, an exact fit was not possible because of two reasons: first there are irreversible reactions, and second, some of the reversible reactions do not satisfy detailed balance. The reaction equations along with their participating species are given in S1 Table. S2 Table and S3 Table compare the parameter amounts of the Kholodenko et al. model with the ones from our reconstructed bond graph model.

The code to convert kinetic parameters into bond graph equivalents for the EGFR pathway is accessible from: https://github.com/Niloofar-Sh/EGFR_MAPK/tree/main/EGF.

#### 2.3.2 MAPK cascade module

Fig 6 shows the schematic of the MAPK cascade. Each oval trajectory in Fig 6.A represents a cycle. The stimulus signal is amplified sequentially through the cycles in the cascade. MKKK is activated through a single phosphorylation phase by a kinase (Ras) and turns into MKKKP [33]. MKK and MK each phosphorylates in two steps and ultimately produce MKKPP and MKPP. The phosphorylated product of each layer plays a kinase role for the phosphorylation phase in the next downstream layer. Simultaneously, an opposing phosphatase dephosphorylates the product of each cycle (shown by backward arrows) [34]. Each layer in Fig 6.A is dephosphorylated by a specific phosphatase: MKKK-Pase in the first layer, MKK-Pase in the second layer, and MK-Pase in the third layer. The dual phosphorylation-dephosphorylation mechanisms in the second and third layers act as amplification, generating ultrasensitive responses.

**Fig 6.**
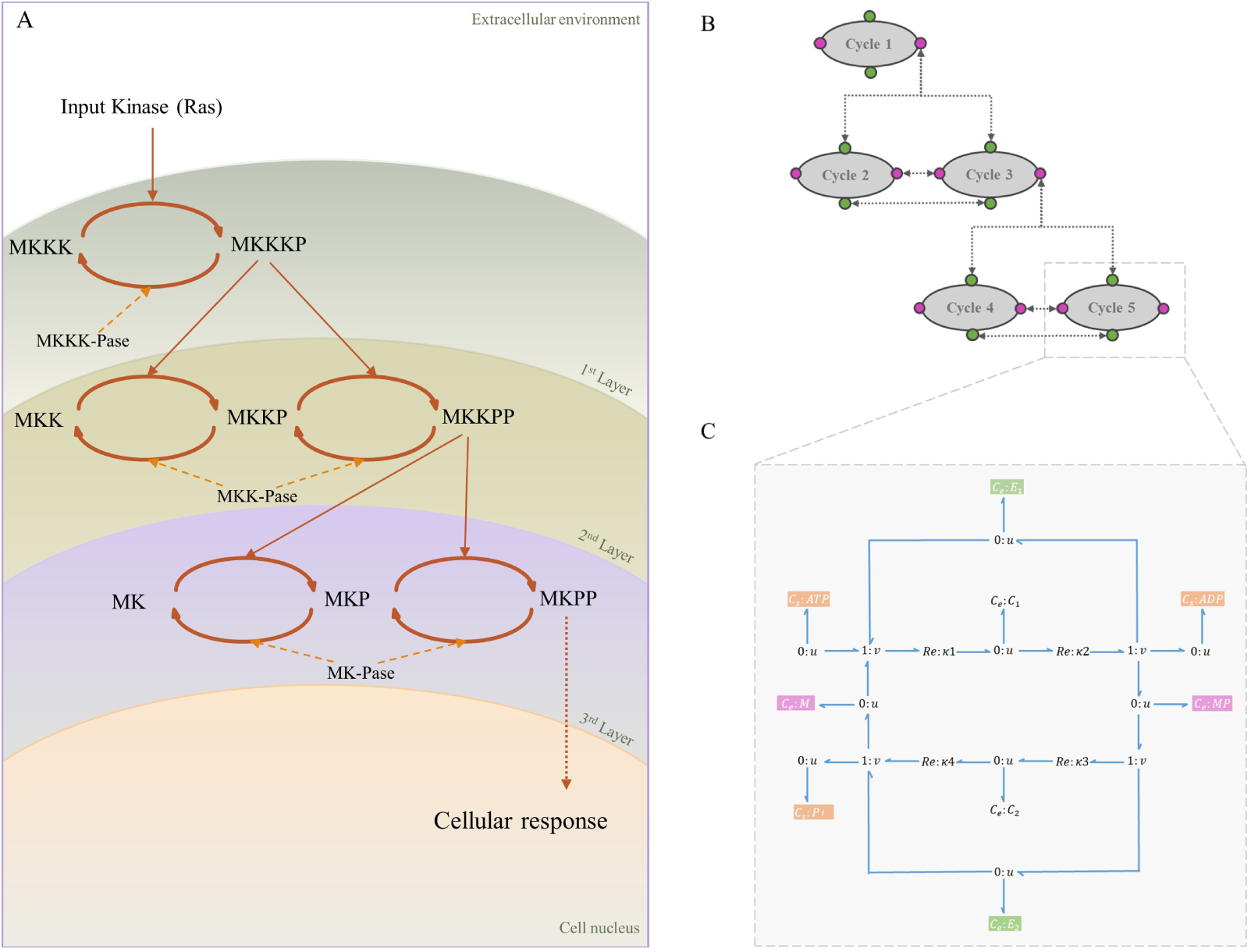
Structure of MAPK cascade. (A) Kinetic representation. The stimulus from the extracellular environment is received (Ras) and transmitted through the MAPK cascade to the cell nucleus. The layers demonstrate the cycles with the same kinase and phosphatase enzymes; (B) MAPK cascade with five modules. Linking species are shown in colours where green corresponds to the linking enzymes and pink corresponds to unphosphorylated/phosphorylated mitogen proteins. Arrows show the links between the modules; (C) The symbolic bond graph model of each cycle. Sources of potential with fixed concentrations (*C_s_*:ATP, *C_s_*:ADP, and *C_s_*:Pi) are shown in orange. (Same sources of potential within the modules are omitted in (A) and (C) for clarity).

Although the species in each cycle are different, the structures of the cycles are the same. The similarity in the structures enables us to break the cascade down into five modules of cycles. Hence, a symbolic bond graph module for a cycle could be created and reused. Fig 6.B shows the modular representation of the cascade by reusing the bond graph module in Fig 6.C. Pan et al. modularized the cascade into phosphorylation/dephosphorylation modules linked by mitogen proteins. We included the mitogen proteins into the symbolic bond graph module to facilitate the composition of the modules. Also, we modelled the MAPK cascade in the absence of feedback. The code for the modular bond graph model of the MAPK cascade in BondGraphTools is accessible from: https://github.com/Niloofar-Sh/EGFR_MAPK/tree/main/MAPK%20cascade.

In the literature, several models of EGFR-Ras-MAPK signalling have been developed considering the involvement of MKKP as an enzyme in phosphorylation of MK and MKP [32] as well as negative feedback from MKPP to the upstream EGFR pathway [33–37]. This paper aims to demonstrate the reusability and composition of bond graph modules; thus, further involvements and feedback loops are not considered in our composition procedure. However, to verify the behaviour of the final composed model, we studied the response of our bond graph EGFR-Ras-MAPK model under the condition of adding negative feedback from MKPP to incorporate as an inactivating enzyme in the first cycle as studied by Kholodenko et al. [35].

In the next section, we illustrate the workflow by applying it to an example biochemical network: EGFR-Ras-MAPK signalling pathway.

### 2.4 Applying the composition method to EGFR-Ras-MAPK signalling pathway

Here, we implement our model composition method (described in Section 2.1.2) to generate a model of EGFR-Ras-MAPK signalling pathway. The EGFR-Ras-MAPK signalling pathway is comprised of two major modules: the EGFR pathway and MAPK cascade. Since MAPK cascade includes five structurally repetitive cycles, we broke it down into five sub-modules. As such, we need two template bond graph modules; one for the EGFR pathway (Fig 5) and one for the cycles in MAPK cascade (Fig 6.C). The connectivity matrix for each module in *csv* format and the annotated CellML files for the parameters of each module/sub-module are available on GitHub: https://github.com/Niloofar-Sh/EGFR_MAPK. Due to the limited size of uploaded files on GitHub, the required reference ontologies for the current model composition (CHEBI, FMA, OPB, and GO) are not provided in the repository (refer to Section 2.1.1 for ontologies).

As well as checking for inconsistencies among the values of similarly annotated components and parameters, components will go through the composition process. In the composition process, if they are mergeable, only one is kept and the rest will be removed from the modules (the list of components for each module will be updated). Furthermore, the rows and columns of the connectivity matrices that correspond to the removed components will be deleted.

Ultimately, a connectivity matrix describing all the connections between the components of the composed network is needed. This is done by integrating the connectivity matrices of all the modules into one. All the connectivity matrices are put consecutively in the diagonal direction of a zero square matrix. Thus, the number of rows/columns equals the total number of components in the system. Subsequently, where we need a bond between two modules, an additional 1 will be inserted in the matrix. An example demonstrating this is shown in Fig 7.

**Fig 7.**
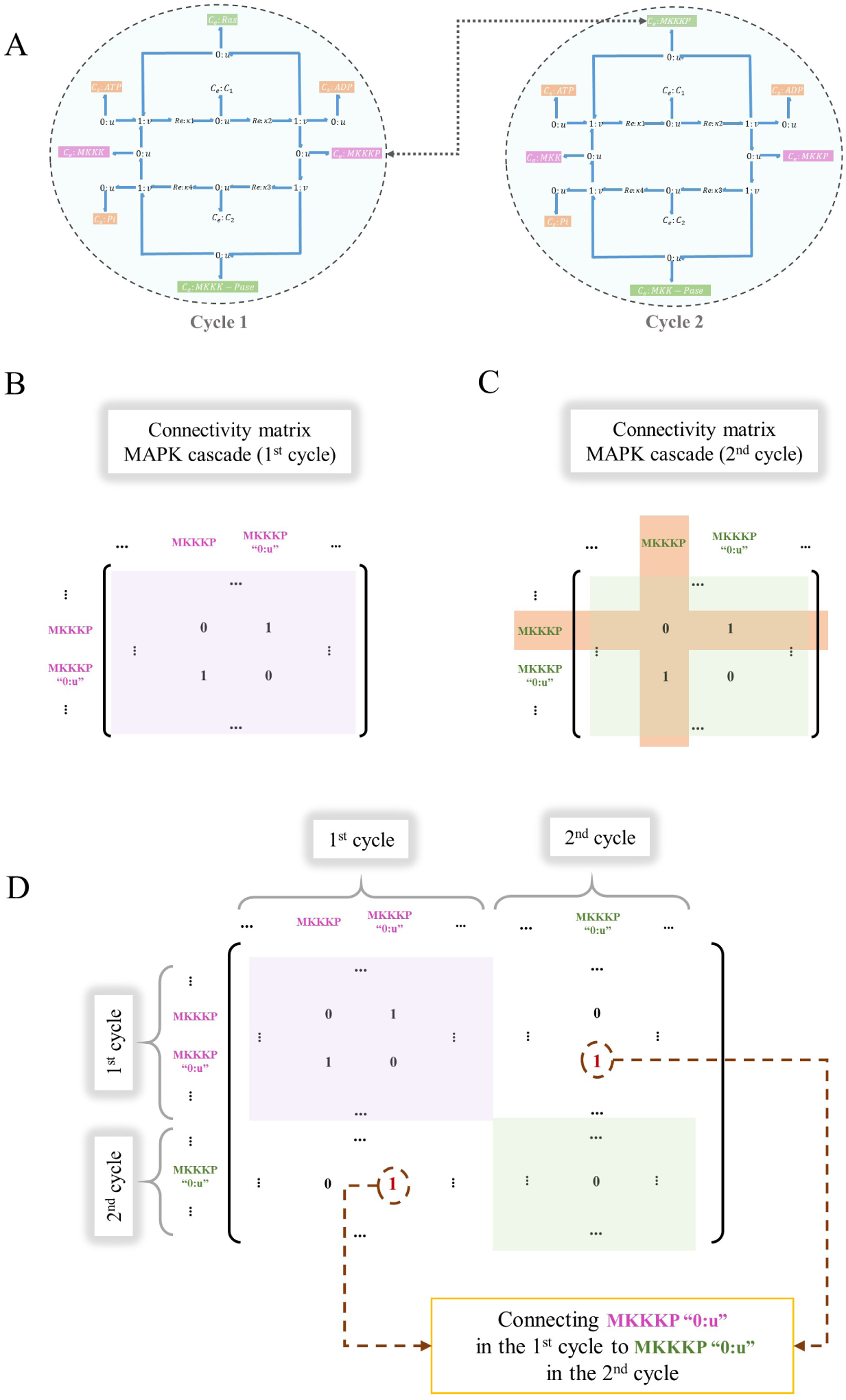
Construction of the whole-system connectivity matrix for a composed model. The procedure is illustrated by integrating two connectivity matrices (1*^st^* and 2*^nd^* cycles in MAPK cascade). Initially, the two cycles had identical connectivity matrices. (A) MKKKP is a common component between the first and second cycle; (B) The connectivity matrix for the 1*^st^* cycle; (C) The modified connectivity matrix for the 2*^nd^* cycle where the row and column for the common component (MKKKP) will be removed; (D) The placement of the connectivity matrices for each module on the diagonal of the whole-system connectivity matrix. The pink and green boxes indicate the connectivity matrices for the 1*^st^* and 2*^nd^* cycles, respectively. The corresponding ‘0:u’ junctions for MKKKP in the two cycles are connected by inserting two 1s (in red) to represent a bond between them (bidirectional connections between the components require the matrix be symmetric).

### 2.5 Verification

To verify the behaviour of the composed EGFR-Ras-MAPK pathway model, we first compare our bond graph estimation of the EGFR module to the original model using the concentration of species. To compare the simulation results between the two models, the normalised root mean square error (NRMSE) was computed as in Eq 7, where *x*^_i_ corresponds to the simulation points of our bond graph estimation, and *x*_i_ corresponds to the Kholodenko et al. model. The normalisation was performed relative to the difference of maximum and minimum data of the reference model in each simulation.

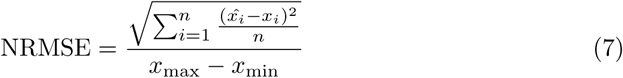

Second, we studied the steady-state behaviour of the phosphorylated kinases at the terminal level of each layer of the MAPK cascade under varying stimulus strengths.

This denoted how we should expect the kinases to respond to any stimulus coming from the upstream levels (here, the RShGS complex).

To further study our composed model, we observed its behaviour under two more conditions: a) We added negative feedback from the terminal phosphorylated kinase in the last layer of the cascade (MKPP) to the dephosphorylation reaction in the initial layer [35] (Fig 8). The effect of adding the negative feedback was then observed and qualitatively verified. b) We simulated the model for different intracellular ATP concentrations and monitored how the concentration of activated kinases was correlated to this change.

**Fig 8.**
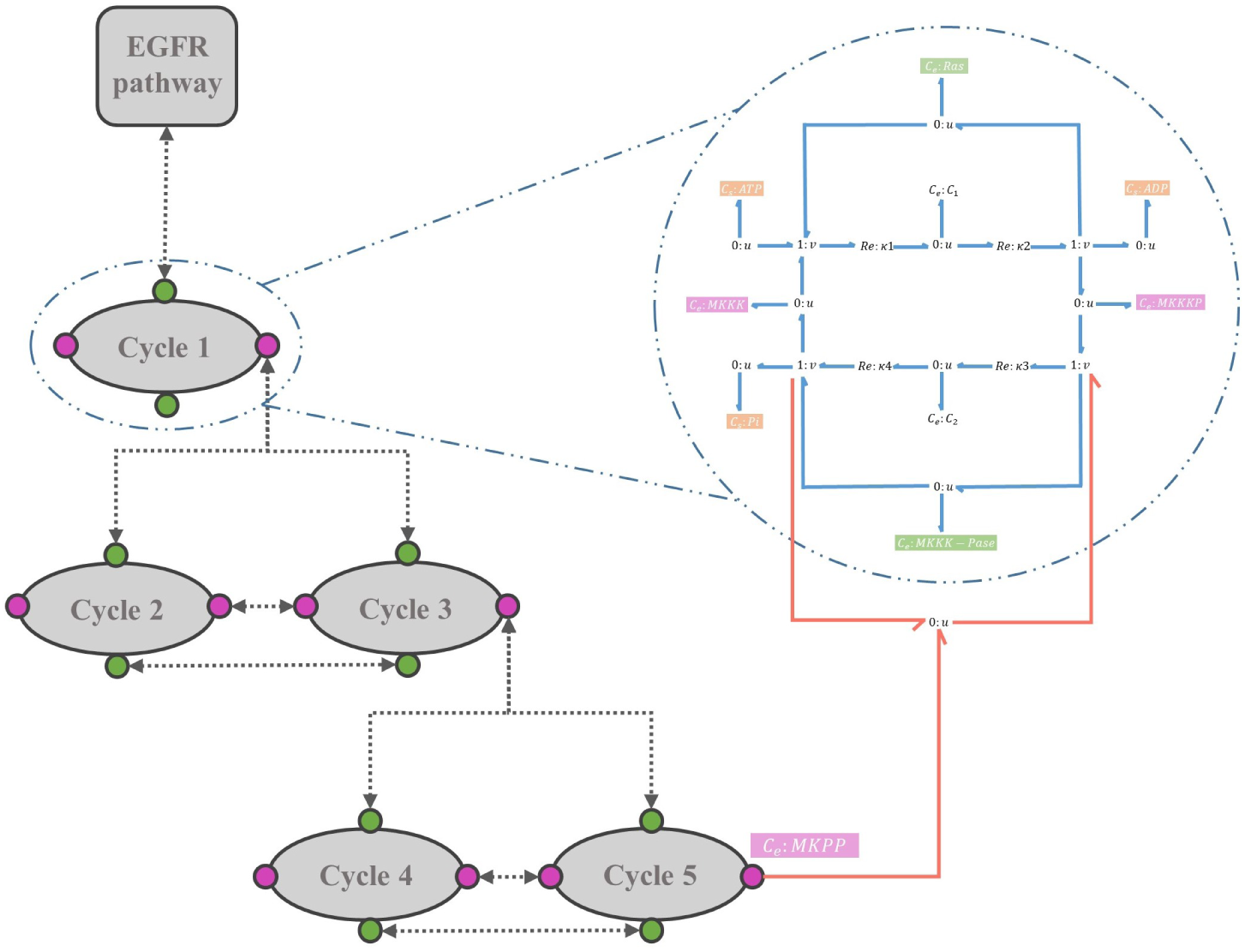
Bond graph schematic of adding negative feedback in the composed EGFR-Ras-MAPK pathway model. The negative feedback loop (red bonds) initiates from MKPP and has an enzymatic role in the first layer’s dephosphorylation reaction.

## 3 Results

We used our method to merge the modules within the MAPK cascade and between the EGFR and MAPK models, yielding the bond graph configuration of the EGFR-Ras-MAPK signalling pathway. Fig 9 shows how the EGFR and MAPK modules are manipulated to deal with same components existing within the modules. Here, Ras and all ATP-ADP-Pi trio components in the MAPK model are removed while RShGS and all ATP-ADP-Pi trios in the EGFR model are kept and bonded to the ‘0 : *u*’ junctions corresponding to the removed components.

**Fig 9.**
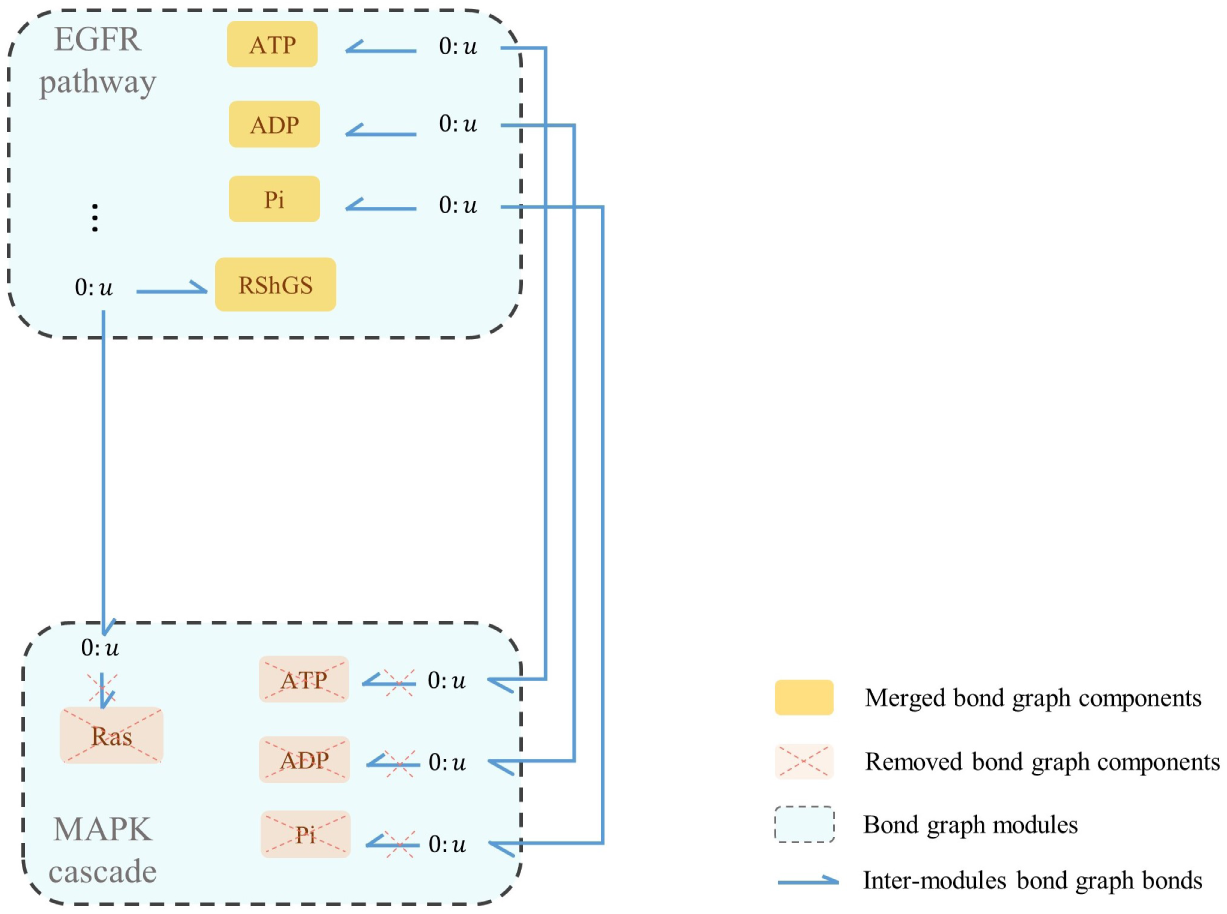
The composed modular bond graph model of EGFR-Ras-MAPK signalling pathway. The blue dashed boxes represent the bond graph modules, the yellow boxes show the merged common components between the modules (each sharing a common potential by a ‘0 : *u*’ junction), and the blue harpoons represent the bonds between the modules and common components. The inter-module bonds, along with the internal bonds between the components in each module, are defined and automatically applied to the model using the whole connectivity matrix. All the six modules also share common potentials with *C_S_* :ATP, *C_S_* :ADP, and *C_S_* :Pi.

To check the function of our composed model, we verified the simulations in two steps: 1. verification of each bond graph module separately (EGFR and MAPK); and 2. verification of the bond graph composed model (EGFR-Ras-MAPK signalling pathway).

### 3.1 Verification of bond graph modules

• **EGFR:** Our approach requires models to be expressed as bond graphs. A bond graph equivalent of the EGFR pathway was not available, which motivated us to convert an existing kinetic model of the EGFR pathway into an equivalent bond graph form. An exact conversion was not possible due to the existence of irreversible reactions and not explicitly accounting for mass conservation. Hence, we approximated the non-bond graph irreversible reactions with bond graph equivalents and included the missing metabolites ATP, ADP, and Pi to provide the energy required to approximate the irreversible reactions.

The conversion of the kinetic EGFR model into bond graphs was performed by solving a linear matrix of equations for the constraints. The species’ responses in the EGFR bond graph module were observed and compared to the ones derived from the Kholodenko et al. model. The responses of four exemplar species in the pathway are demonstrated in Fig 10, and the NRMSE is computed for each comparison in percentage. We see that the bond graph equivalent of the EGFR module functioned similarly to the original kinetic model, although the equations could not be solved perfectly. This implies that the kinetic parameters of the Kholodenko et al. model are not thermodynamically consistent. The bond graph equivalent represents a close-match approximation of the original model in a thermodynamically consistent manner.

**Fig 10.**
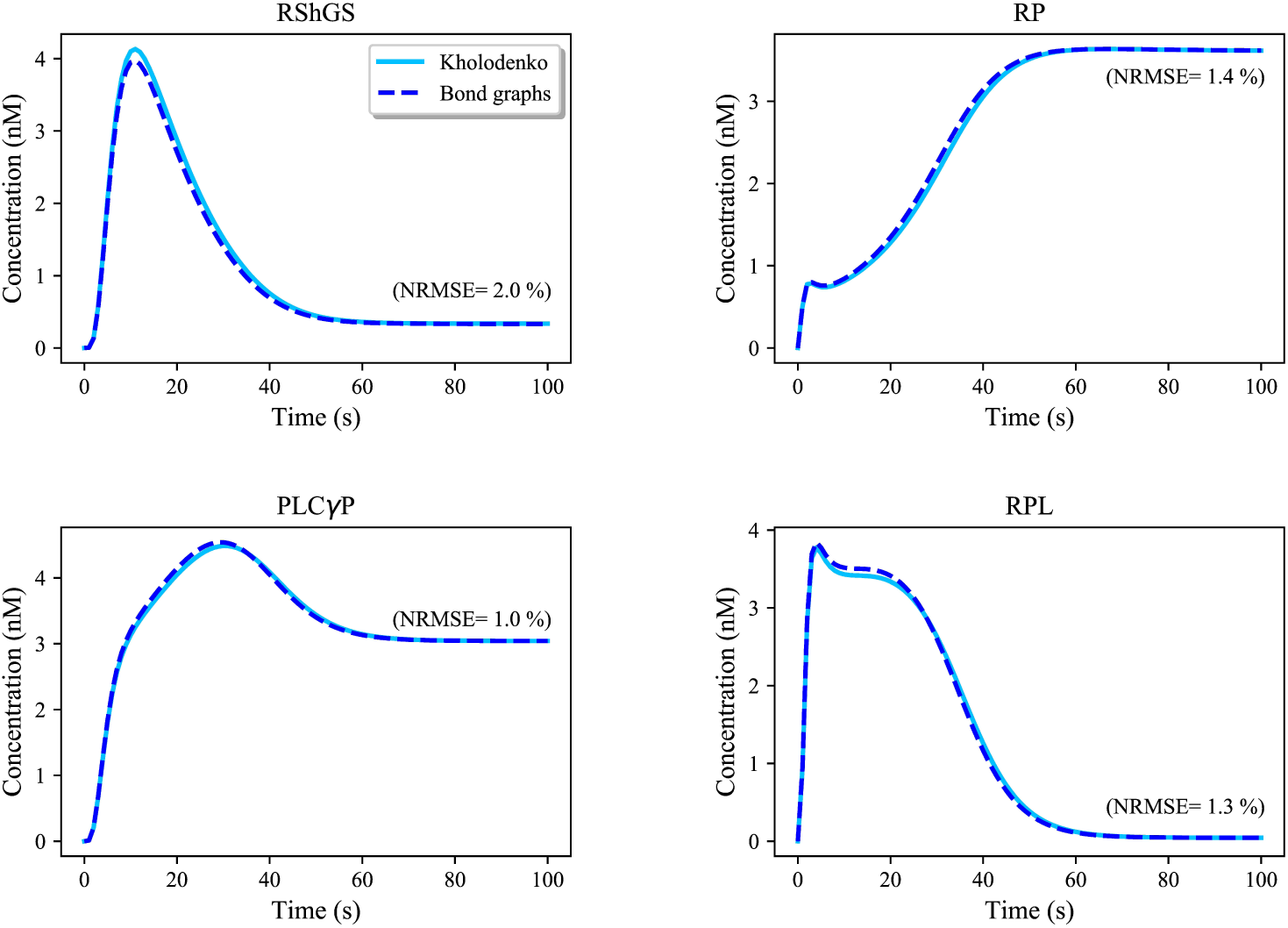
Comparison between the Kholodenko et al. EGFR model and its bond graph approximation. The simulations are given for four exemplar species in the pathway. NRMSE is calculated for each comparison in percentage. The initial concentration of EGF (the initiative molecule in the EGFR module) was 680 nM.

• **MAPK:** The bond graph model of the MAPK cascade was developed by Pan et al. [21]. We have reused the model here with slightly different configuration of the modules.

The bond graph version of MAPK cascade in BondGraphTools was simulated with an initial amount of Ras = 3 *×* 10*^−^*^5^ (*µ*M). A minor increase in the concentration of the input kinase results in amplified sigmoidal responses of downstream kinases, referred to as ultrasensitivity (S1 Fig) [35]. Amplification in the layers of MAPK cascade form the ultrasensitive responses, *i.e.*, single phosphorylation-dephosphorylation in the first layer and dual phosphorylation-dephosphorylation in the second and third layers. At this point, we plotted the steady-state responses of the activated kinases against a range of input concentrations (10*^−^*^8^ – 10^0^ (*µ*M)) in Fig 11. Note how for inputs less than 7 *×* 10*^−^*^5^ (*µ*M) MKPP reaches a higher concentration than MKKPP while MKKPP overtakes MKPP for higher input concentrations. S2 Fig shows the relative activation of the kinases indicating the lower the layer in the MAPK cascade, the smaller the input concentration activates the kinases [21]. Note that the MKPP (third layer) activation curve is steeper compared to MKKPP (second layer) and MKKKP (first layer), projecting that a higher increase in the stimulus is required for MKKKP to reach its maximum response compared to MKKPP and MKPP (Table 1). The analysis of the behaviour of the MAPK module assisted us to predict how the kinases will respond to the input kinase (Ras) coming from the upstream module (EGFR pathway) and validate our composed bond graph model.

**Fig 11.**
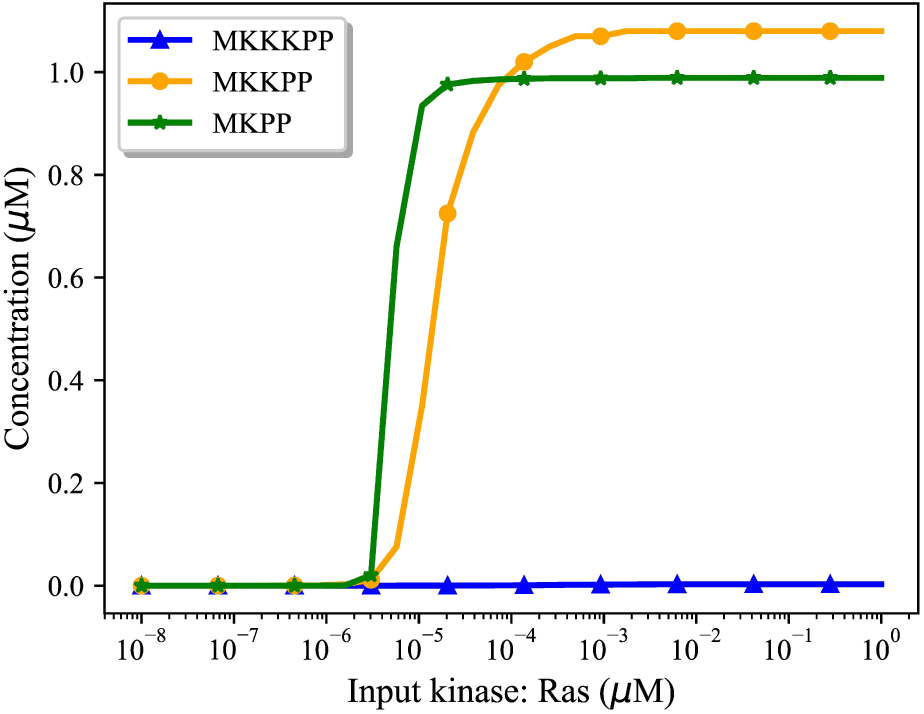
The steady-state responses of the activated kinases for different input amounts. The input Ras concentration is expressed on a logarithmic scale.

**Table 1.**
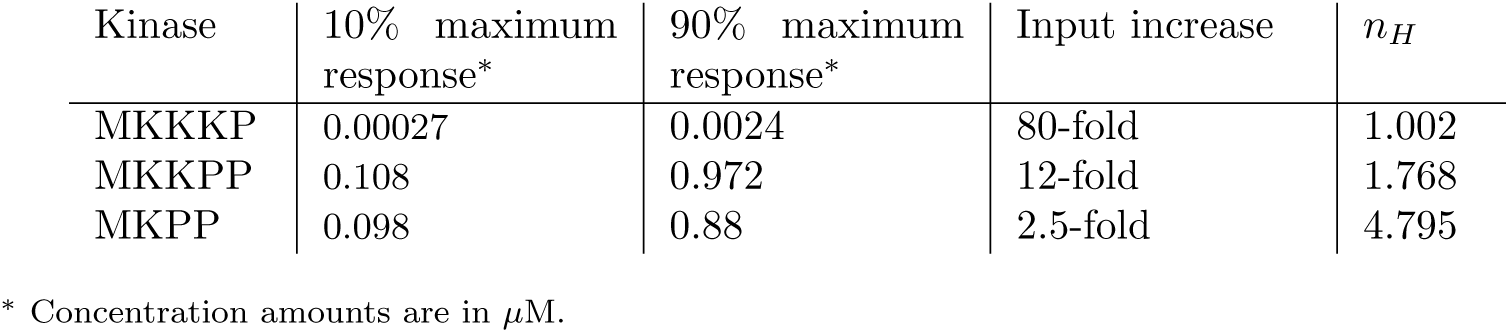
Input differences in reaching from 10% to 90% of maximum concentration in kinases.

| Kinase | 10% maximum response* | 90% maximum response* | Input increase | $n_H$ |
| --- | --- | --- | --- | --- |
| MKKKP | 0.00027 | 0.0024 | 80-fold | 1.002 |
| MKKPP | 0.108 | 0.972 | 12-fold | 1.768 |
| MKPP | 0.098 | 0.88 | 2.5-fold | 4.795 |

\* Concentration amounts are in $\mu\text{M}$ .

Table 1 delineates the required stimulus increase for each activated kinase to reach from 10% of its ultimate concentration to 90%. This affirms the ultrasensitive responses to the input as we go to the lower layers of the cascade. To estimate the ultrasensitivity in sigmoidal input-output curves, the Hill coefficient (nH) is also calculated per activated kinase as per Eq 8, where EC90 and EC10 are the input values required to produce 90% and 10% of the maximal response, respectively [38]. The greater the Hill coefficient than 1, the smaller input value is required for the concentration transition from 10% to 90% of its maximum amount. The figures are consistent to the predicted Hill coefficients for MAPK cascade in work by Huang & Ferrell [39].

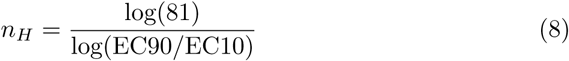

### 3.2 Verification of the bond graph composed EGFR-Ras-MAPK model

We investigated the behaviour of our bond graph composed model (EGFR-Ras-MAPK pathway) under three conditions: **without negative feedback**, **with negative feedback**, and **different ATP concentrations** to examine the functionality of our model under varying conditions. Each of these three conditions imply qualitatively predictable changes in the behaviour of the whole network which we aim to investigate in our composed model.

#### • Without negative feedback

The simulated time courses of the three activated kinases (MKKKP, MKKPP, and MKPP) in the composed bond graph model of the EGFR-Ras-MAPK pathway are shown in Fig 12.A. Fig 12.B predicts the activated kinases at steady-state for various input concentrations. The semi-steady-state concentration of the input kinase (Ras) was 14.18 nM, which is indicated by the purple dashed line in Fig 12.B. The intersection of this line with the MKKKP, MKKPP, and MKPP concentrations shows the expected steady-state concentrations of the aforementioned kinases.

**Fig 12.**
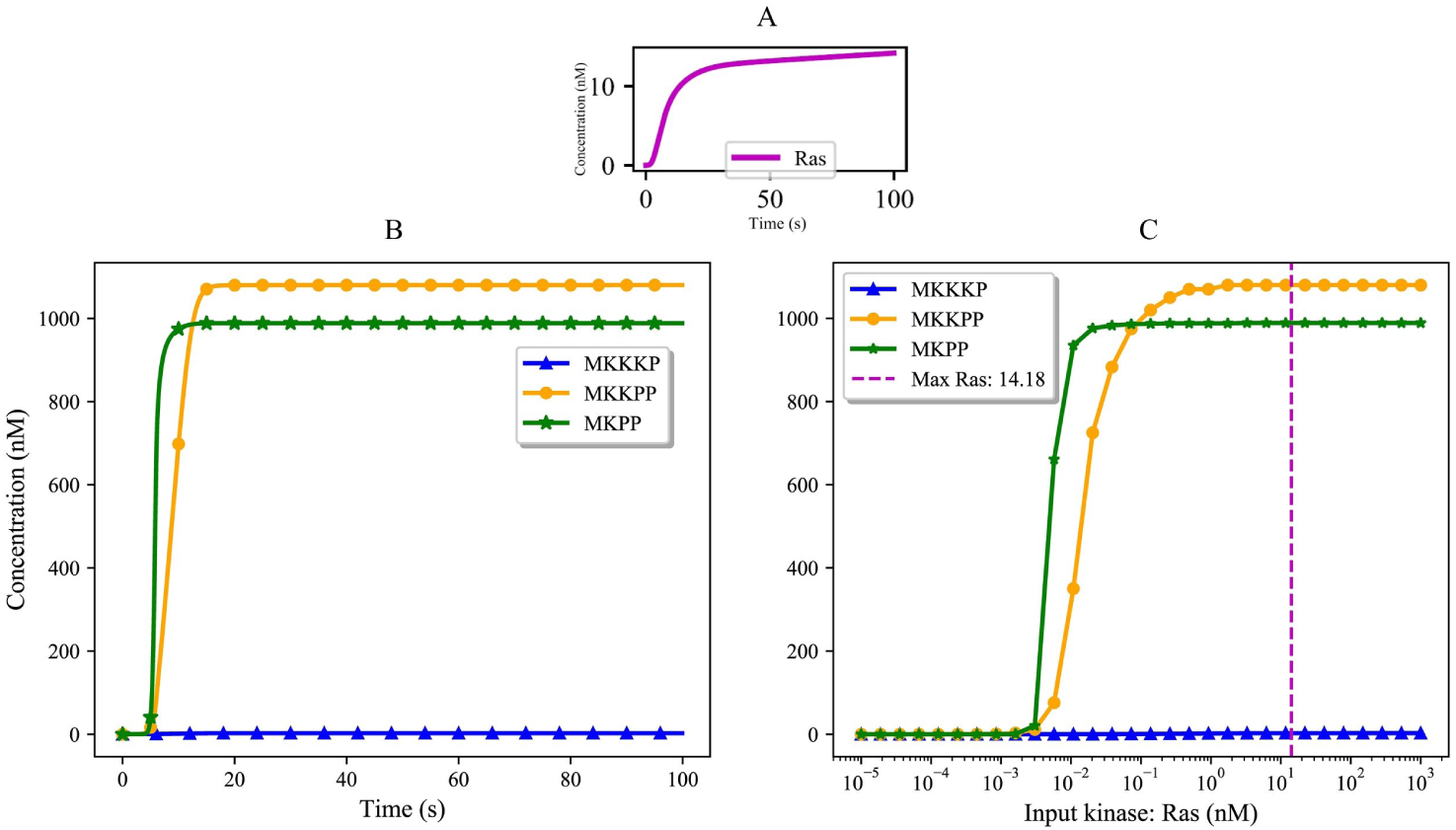
Responses of activated kinases to the input kinase (Ras) in the composed bond graph model of the EGFR-Ras-MAPK pathway. (A) Ras signal; (B) Ultrasensitivity in the composed bond graph models of the EGFR-Ras-MAPK pathway; (C) Predicted steady-state activation of the kinases. The purple dashed line shows the steady-state imposed Ras signal.

#### • With negative feedback

Negative feedback in MAPK cascade may lead to inhibited responses or oscillations depending on the stability points of the system [37]. The activated kinases respond differently when a negative feedback loop is added to the system. This feature was also explored in our composed bond graph model of the EGFR-Ras-MAPK pathway.

Fig 13 compares the steady-state responses of terminal kinases in the MAPK cascade model in two cases: without negative feedback (Fig 13.A) and with negative feedback (Fig 13.B). Under the effect of a negative feedback loop in the MAPK cascade the kinases require higher input Ras concentration to reach their steady-state concentrations. The right shift to the curves in Fig 13.B shows the expected functionality of the MAPK cascade module in the presence of negative feedback along with reduced amplification.

**Fig 13.**
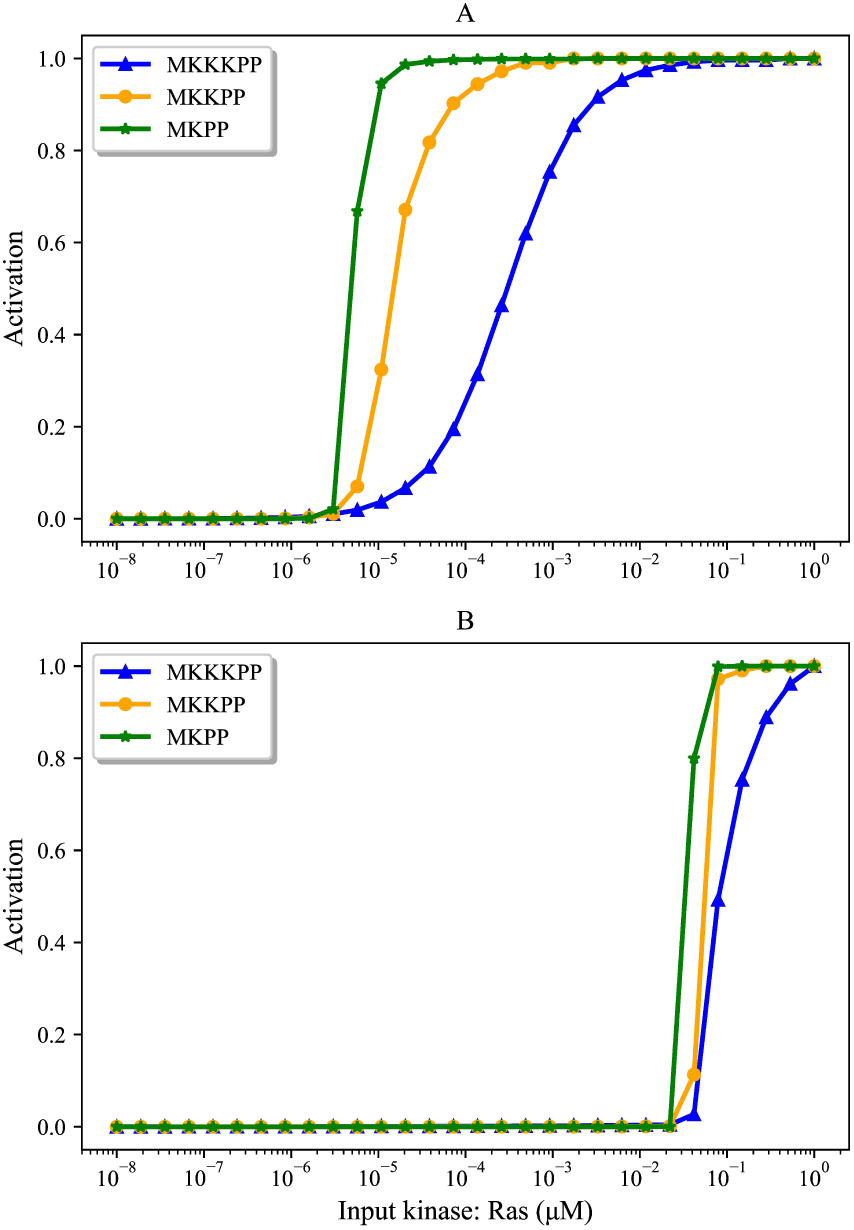
Activation of terminal kinases with and without negative feedback. (A) Without negative feedback; (B) With negative feedback. The input kinase (Ras) concentration is expressed on a logarithmic scale

Next, we show how the terminal kinases respond in our bond graph composed model of EGFR-Ras-MAPK pathway in the presence of the same negative feedback loop. Fig 14 shows the inhibited responses of the terminal kinases and subsequently, the significant delay in reaching their steady-states (compare with Fig 12.B). The added Negative feedback in EGFR-Ras-MAPK model strengthens the dephosphorylation reaction in the first layer of the MAPK cascade module which receives the Ras stimulus. This strengthened dephosphorylation inhibits its corresponding phosphorylation pair and affects phosphorylation in all the proceeding layers.

**Fig 14.**
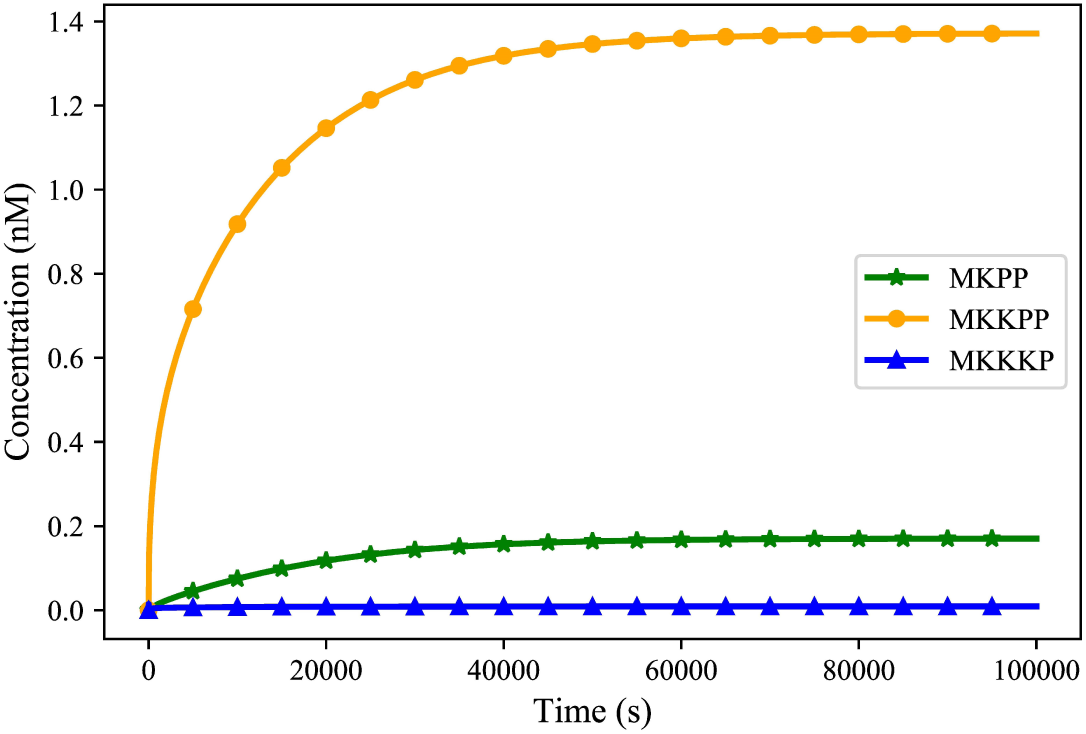
Time course behaviour of the terminal kinases in the EGFR-Ras-MAPK bond graph model with negative feedback.

#### • ATP concentration

ATP is one of the species involved in prompting wound responses that activates MAPK pathways in cells [40]. As such, ATP shortage causes delays or failure in activating kinases, and as a result, dysfunction in wound healing responses. The production of ATP in cells might be blocked or reduced due to multiple reasons, such as mitochondrial disorders, ageing, or very intense exercises [41–43].

The impact of ATP concentration on the behaviour of the bond graph EGFR-Ras-MAPK model was investigated by clamping the ATP concentration at **15%**, **35%**, **81%**, and **100%** of its baseline level (Fig 15). Fig 15.A-C illustrate how different levels of cellular ATP (energy) influence the behaviour of activated kinases and also confirm that ATP shortage induces a delay in the responses. Fig 15.D compares the steady-state concentration of MKKKP, MKKPP, and MKPP against various ATP concentrations relatively. The lower the ATP production, the lower the steady-state concentration of MKKKP, MKKPP, and MKPP, highlighting the importance of energy for the function of the pathway.

**Fig 15.**
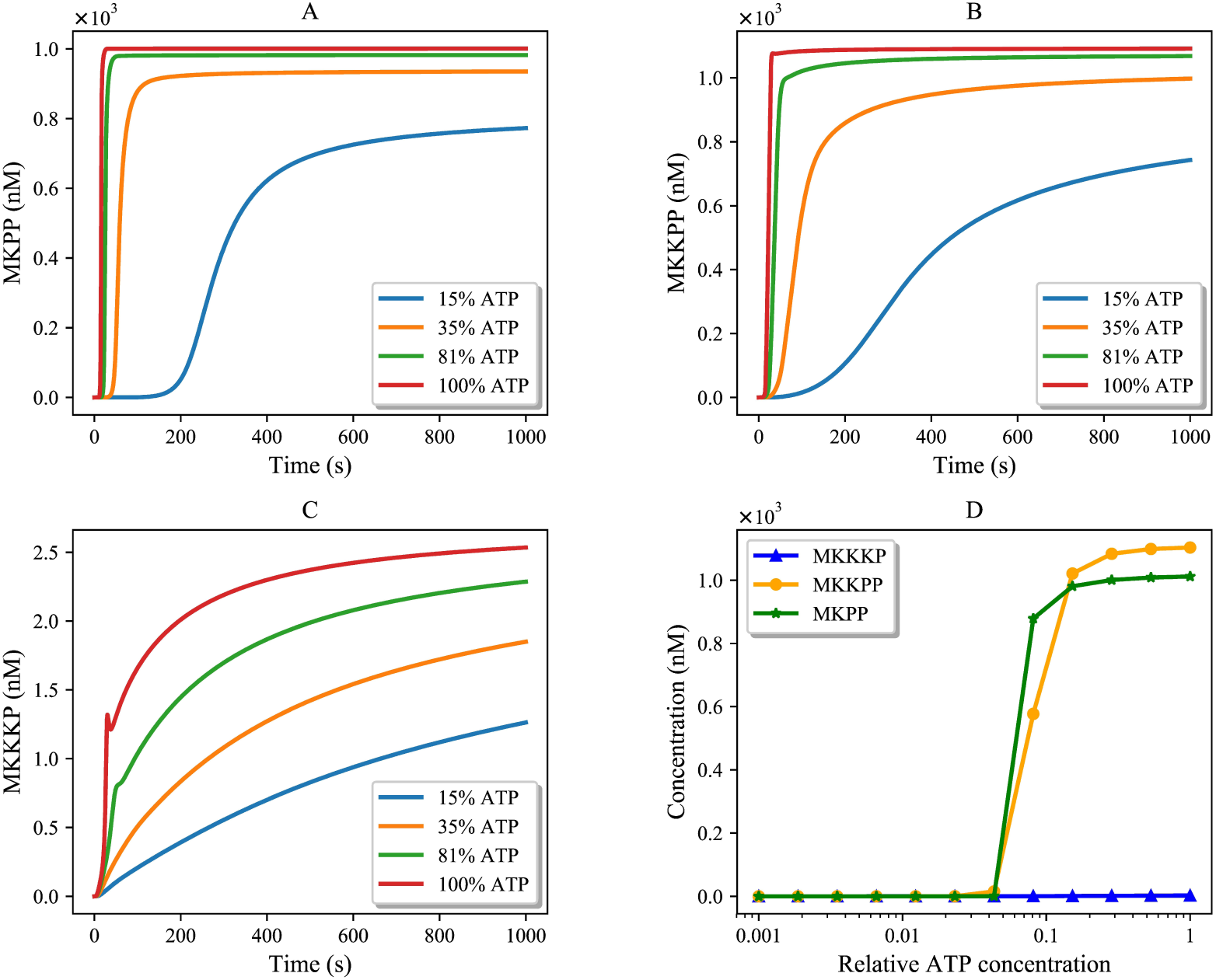
Effect of different levels of ATP concentration on activated kinases. (A) MKPP; (B) MKKPP; (C) MKKKP; (D) Steady-state concentration of MKKKP, MKKPP, and MKPP against relative ATP concentration.

## 4 Discussion

In this paper, we introduced a generic approach to assemble computational models in biology without starting from scratch. This was enabled by constructing symbolic bond graph modules of biophysical systems and obtaining the required parameters from existing models. To extract and allocate the parameters, the conventional target model needs to be fully and properly annotated. Here, we have selected models encoded in CellML, but the approach can be applied to models in other formats, such as SBML, as long as they can include a semantic description of the system being modelled. For biochemical reactions, if the parameters are thermodynamically inconsistent, they are converted into bond graph compatible ones. The modules will then automatically combine when the common components (species) among them are merged. The resulting composed bond graph model complies with the laws of physics and can be coupled to other bond graph modules. As an example, we applied our method to the EGFR-Ras-MAPK pathway.

Our composed model of EGFR-Ras-MAPK signalling pathway is different from the ones in the literature in three ways which prevents us from conducting a direct comparison:

- Our reference models of EGFR pathway and MAPK cascade are adopted from different sources which included/excluded some reactions or feedback effects;
- Some reactions in the original EGFR model were irreversible and therefore thermodynamically infeasible;
- RShGS and RGS both trigger the activation of Ras protein through an intermediate molecule that serves as a switch for the transmission of the signal to the downstream MAPK cascade and yields Ras. Since none of our reference models included this intermediate and we could only merge one species with another one, we selected RShGS to be merged with the Ras protein in the MAPK cascade module due to its prominent role in localising Ras compared to RGS [44, 45].

Thus, we validated the composed model by comparing the behaviour of the terminal kinases to the predicted behaviours from running the MAPK cascade model solely (Fig 11). The purple dashed line in Fig 12.C denotes the steady-state concentration forecast of each activated kinase at Ras = 14.18 (nmol). The results gained from our composed model in Fig 12.B comply with the predicted ones in Fig 12.C.

Merging components across models might raise inconsistencies in their parameters. Here, RShGS in the EGFR model was merged with Ras in the MAPK model. These two species have different initial values and thermodynamic constants in their corresponding models. In such cases, our framework flags inconsistent values for same species. This is solved by asking the user to either select one of the values or insert a new value for the flagged parameter. Since the user may not have the relevant expertise, we aim to provide users with an evaluation of the ambiguous parameter in multiple models available on PMR in the future. This will give the user a better awareness of the range of values for uncertain parameters.

As an improvement to our previous approach [17], the present framework overcame the aforementioned limitations:

1. No mathematical formulation of bond graphs is required in the CellML modules (formulating symbolic bond graph modules in BondGraphTools is more straightforward and less error-prone);
2. Auxiliary variables are not needed in the CellML modules as linking ports (ports are automatically detected using ‘white box’ approach by finding similar annotations);
3. Instead of the semi-automated SemGen merger tool, our approach integrates the modules in a fully-automated manner (our implementation automatically merges the modules and performs the required structural changes).

Currently, our model composition approach is capable of detecting exact matches. However, this approach could be improved by allowing the user to specify mergeable components from a shortlist of similarly annotated ones. In the future, if the scientific community defines a globally accepted standard to unify the annotation of similar biological models, finding matching metadata among the models will be facilitated.

Our energy-based model composition approach is designed to link mathematical models encoded in CellML to their bond graph equivalent and compose them in a consistent and physics-based environment. Currently, there is no general method of automatically converting mathematical models into bond graphs, and each model requires domain-specific expertise to generate a similar bond graph form. To reuse and compose the massive number of existing biological models, the community should either push the researchers to build thermodynamically consistent and physically plausible models or encourage the researchers to develop computational tools that convert existing biological models into bond graphs.

If a model follows the laws of physics and thermodynamics, it can be directly converted into bond graphs. Otherwise one must make assumptions to produce a bond graph that approximates the original model. To facilitate such decisions, we propose establishing an evaluation system to check whether the original model is physically realistic or not. If the model cannot represent a physically plausible system and its bond graph approximation does not fit the data, it highlights some inconsistencies in the original model that must be noted and fixed.

The ultimate goal of applying our model composition method is to provide a foundation for future tool developments to convert any arbitrary CellML model into bond graphs and then convert it back to a CellML file. We require the bond graph conversion for appending, deleting, and editing modules. This allows us to firstly avoid any errors or confusions during the process, and secondly, make sure that the model conserves energy and mass and remains thermodynamically and physically consistent as we modify it. Eventually, the generated mathematical equations in the bond graph environment can be converted into CellML for simulation and reproducibility.

Models are constructed in different units for parameters and various scales of amounts. Coupling arbitrary models will alter their boundary conditions which induces differences that propagate throughout the models. In the future, we plan to apply nondimensionalization to remove dependencies to the measured units across the models and generate unified composed models, regardless of their units [46].

Nondimensionalization is especially useful in models that are described by differential equations. In this systematic technique, all variables and parameters become unitless by rescaling them relative to a reference value.

## 5 Conclusion

We have developed a method that automates the integration of biosimulation models. We utilised the SemGen annotator tool to add metadata to CellML models and the Python library BondGraphTools to generate the bond graph template of models. Describing the bonds between bond graph components with connectivity matrices helped us conveniently delete or add bonds/components to the modules. This minimises user error when a structural change is required in complex systems. Here we have presented a method that automates the composition by taking advantage of semantics in the modules and the systematic structural modification using connectivity matrices. We demonstrated the functionality of our method by coupling two biosimulation models and their sub-models. Likewise, several annotated biosimulation models can be integrated automatically if they have common elements/species. This is particularly pivotal when dealing with complex and large biological systems where mathematically merging models requires time-consuming and error-prone post-merging adjustments.

We believe that our method is one of the initial steps toward multiscale cell-to-organ-level model integration.

## 6 Supporting information

**S1 Fig. Ultrasensitivity in MAPK cascade.** For an input kinase of Ras=3 *×* 10*^−^*^5^ (*µ*M), the concentration changes of the activated kinases (MKKKP, MKKPP, and MKPP) show the signal is amplified through each layer.

**S2 Fig. The normalised activation of kinases in the MAPK cascade module for different input amounts (Ras).**

**S1 Table. Reactant(s) and product(s) of each step in EGFR pathway and the reaction rate equations.** Steps 4, 8, and 16 are irreversible reactions, which are approximated by mass action kinetics. *κ_i_*(*i ∈ {*Step*}*) in the reaction rate equations represent the reaction rate constants, *K_x_* (*x ∈ {*Reactants, Products*}*) is the thermodynamic constant of each species, and *q_x_* (*x ∈ {*Reactants, Products*}*) is the concentration amount of each species.

**S2 Table. Original and modified parameters of the species in the EGFR pathway model.**

**S3 Table. Original and modified parameters of the reactions in the EGFR pathway model.**

## Acknowledgments

NS would like to thank Yuda Munarko for his helpful comments and suggestions.

## References

1. Carrera J, Covert MW. Why build whole-cell models? Trends in cell biology. 2015;25(12):719–722.

2. Cooling MT, Nickerson DP, Nielsen PM, Hunter PJ. Modular modelling with Physiome standards. The Journal of physiology. 2016;594(23):6817–6831.

3. Yu T, Lloyd CM, Nickerson DP, Cooling MT, Miller AK, Garny A, et al. The physiome model repository 2. Bioinformatics. 2011;27(5):743–744.

4. Le Novere N, Bornstein B, Broicher A, Courtot M, Donizelli M, Dharuri H, et al. BioModels Database: a free, centralized database of curated, published, quantitative kinetic models of biochemical and cellular systems. Nucleic acids research. 2006;34(suppl 1):D689–D691.

5. Clerx M, Cooling MT, Cooper J, Garny A, Moyle K, Nickerson DP, et al. CellML 2.0. Journal of Integrative Bioinformatics. 2020;17(2-3).

6. Hucka M, Finney A, Sauro HM, Bolouri H, Doyle JC, Kitano H, et al. The systems biology markup language (SBML): a medium for representation and exchange of biochemical network models. Bioinformatics. 2003;19(4):524–531.

7. Malik-Sheriff RS, Glont M, Nguyen TV, Tiwari K, Roberts MG, Xavier A, et al. BioModels—15 years of sharing computational models in life science. Nucleic acids research. 2020;48(D1):D407–D415.

8. Smith LP, Hucka M, Hoops S, Finney A, Ginkel M, Myers CJ, et al. SBML Level 3 package: Hierarchical Model Composition, Version 1 Release 3. Journal of Integrative Bioinformatics. 2015;12:603 – 659.

9. de Bono B, Safaei S, Grenon P, Hunter PJ. Meeting the multiscale challenge: representing physiology processes over ApiNATOMY circuits using bond graphs. Interface Focus. 2017;8.

10. Gawthrop PJ, Crampin EJ. Energy-based analysis of biochemical cycles using bond graphs. Proceedings of the Royal Society A: Mathematical, Physical and Engineering Sciences. 2014;470(2171):20140459.

11. Paynter H. Analysis and Design of Engineering Systems/Paynter HM;.

12. Oster G, Perelson A, Katchalsky A. Network thermodynamics. Nature. 1971;234(5329):393–399.

13. Oster GF, Perelson AS, Katchalsky A. Network thermodynamics: dynamic modelling of biophysical systems. Quarterly reviews of Biophysics. 1973;6(1):1–134.

14. Cellier F. Modeling Chemical Reaction Kinetics. In: Continuous System Modeling. Springer, New York, NY; 1991.

15. Gawthrop PJ, Cursons J, Crampin EJ. Hierarchical bond graph modelling of biochemical networks. Proceedings of the Royal Society A: Mathematical, Physical and Engineering Sciences. 2015;471(2184):20150642.

16. Schamai W, Buffoni L, Fritzson PA. An Approach to Automated Model Composition Illustrated in the Context of Design Verification. Modeling Identification and Control. 2014;35:79–91.

17. Shahidi N, Pan M, Safaei S, Tran K, Crampin EJ, Nickerson DP. Hierarchical semantic composition of biosimulation models using bond graphs. PLoS computational biology. 2021 May 13;17(5):e1008859.

18. Neal ML, Thompson CT, Kim KG, James RC, Cook DL, Carlson BE, et al. SemGen: a tool for semantics-based annotation and composition of biosimulation models. Bioinformatics. 2019;35 9:1600–1602.

19. Cudmore P, Pan M, Gawthrop PJ, Crampin EJ. Analysing and simulating energy-based models in biology using BondGraphTools. bioRxiv. 2021;.

20. Kholodenko BN, Demin OV, Moehren G, Hoek JB. Quantification of Short Term Signaling by the Epidermal Growth Factor Receptor. The Journal of Biological Chemistry. 1999;274:30169–30181.

21. Pan M, Gawthrop PJ, Cursons J, Crampin EJ. Modular assembly of dynamic models in systems biology. PLoS computational biology. 2021 Oct 13;17(10):e1009513.

22. Chapter 3 - Connectivity Matrices and Brain Graphs. In: Fornito A, Zalesky A, Bullmore ET, editors. Fundamentals of Brain Network Analysis. San Diego: Academic Press; 2016. p. 89–113.

23. Neal ML, Cooling MT, Smith LP, Thompson CT, Sauro HM, Carlson BE, et al. A Reappraisal of How to Build Modular, Reusable Models of Biological Systems. PLoS Computational Biology. 2014;10.

24. Gawthrop PJ, Pan M, Crampin EJ. Modular dynamic biomolecular modelling with bond graphs: the unification of stoichiometry, thermodynamics, kinetics and data. Journal of the Royal Society Interface. 2021;18.

25. Wellstead PE. Introduction to physical system modelling. vol. 4. Academic Press London; 1979.

26. Gawthrop P, Smith L. Metamodelling: For bond graphs and dynamic systems. Prentice Hall International (UK) Ltd.; 1996.

27. Borutzky W. Bond graph methodology: development and analysis of multidisciplinary dynamic system models. Springer Science & Business Media; 2009.

28. Pan M, Gawthrop PJ, Tran K, Cursons J, Crampin EJ. A thermodynamic framework for modelling membrane transporters. Journal of theoretical biology. 2018;.

29. Atkins PW, de Paula JC. Physical Chemistry for the Life Sciences; 2005.

30. Gawthrop PJ, Crampin EJ. Bond Graph Representation of Chemical Reaction Networks. IEEE Transactions on NanoBioscience. 2018;17:449–455.

31. Molina J, Adjei A. The Ras/Raf/MAPK pathway. Journal of Thoracic Oncology. 2006;1(1):7–9. doi:10.1097/01243894-200601000-00004.

32. Brightman FA, Fell DA. Differential feedback regulation of the MAPK cascade underlies the quantitative differences in EGF and NGF signalling in PC12 cells. FEBS Letters. 2000;482(3):169–174. doi:https://doi.org/10.1016/S0014-5793(00)02037-8.

33. Sarma U, Ghosh I. Oscillations in MAPK cascade triggered by two distinct designs of coupled positive and negative feedback loops. BMC Research Notes. 2011;5:287 – 287.

34. Orton RJ, Sturm OE, Vyshemirsky V, Calder M, Gilbert DR, Kolch W. Computational modelling of the receptor-tyrosine-kinase-activated MAPK pathway. The Biochemical journal. 2005;392 Pt 2:249–61.

35. Kholodenko BN. Negative feedback and ultrasensitivity can bring about oscillations in the mitogen-activated protein kinase cascades. European journal of biochemistry. 2000;267 6:1583–8.

36. Lake D, Corra S, Muller J. Negative feedback regulation of the ERK1/2 MAPK pathway. Cellular and Molecular Life Sciences. 2016;73. doi:10.1007/s00018-016-2297-8.

37. Arkun Y, Yasemi M. Dynamics and control of the ERK signaling pathway: Sensitivity, bistability, and oscillations. PLoS ONE. 2018;13.

38. Altszyler E, Ventura AC, Colman-Lerner A, Chernomoretz A. Ultrasensitivity in signaling cascades revisited: Linking local and global ultrasensitivity estimations. PLoS ONE. 2017;12.

39. Huang CY, Ferrell JE. Ultrasensitivity in the mitogen-activated protein kinase cascade. Proceedings of the National Academy of Sciences of the United States of America. 1996;93 19:10078–83.

40. Medina-Castellanos E, Esquivel-Naranjo EU, Heil M, Herrera-Estrella A. Extracellular ATP activates MAPK and ROS signaling during injury response in the fungus Trichoderma atroviride. Frontiers in Plant Science. 2014;5.

41. Johnson TA, Jinnah HA, Kamatani N. Shortage of Cellular ATP as a Cause of Diseases and Strategies to Enhance ATP. Frontiers in Pharmacology. 2019;10.

42. Schütt F, Aretz S, Auffarth GU, Kopitz J. Moderately reduced ATP levels promote oxidative stress and debilitate autophagic and phagocytic capacities in human RPE cells. Investigative ophthalmology & visual science. 2012;53 9:5354–61.

43. Hargreaves M, Spriet LL. Skeletal muscle energy metabolism during exercise. Nature Metabolism. 2020; p. 1–12.

44. Sasaoka T, Langlois WJ, Leitner JW, Draznin B, Olefsky JM. The signaling pathway coupling epidermal growth factor receptors to activation of p21ras. The Journal of biological chemistry. 1994;269 51:32621–5.

45. Resat H, Ewald JA, Dixon DA, Wiley HS. An integrated model of epidermal growth factor receptor trafficking and signal transduction. Biophysical journal. 2003;85 2:730–43.

46. Ledder G. Scaling for Dynamical Systems in Biology. Bulletin of Mathematical Biology. 2017;79:2747–2772.

