## Supplementary figures and images for "A semantics, energy-based approach to automate biomodel composition"

### S1 Fig

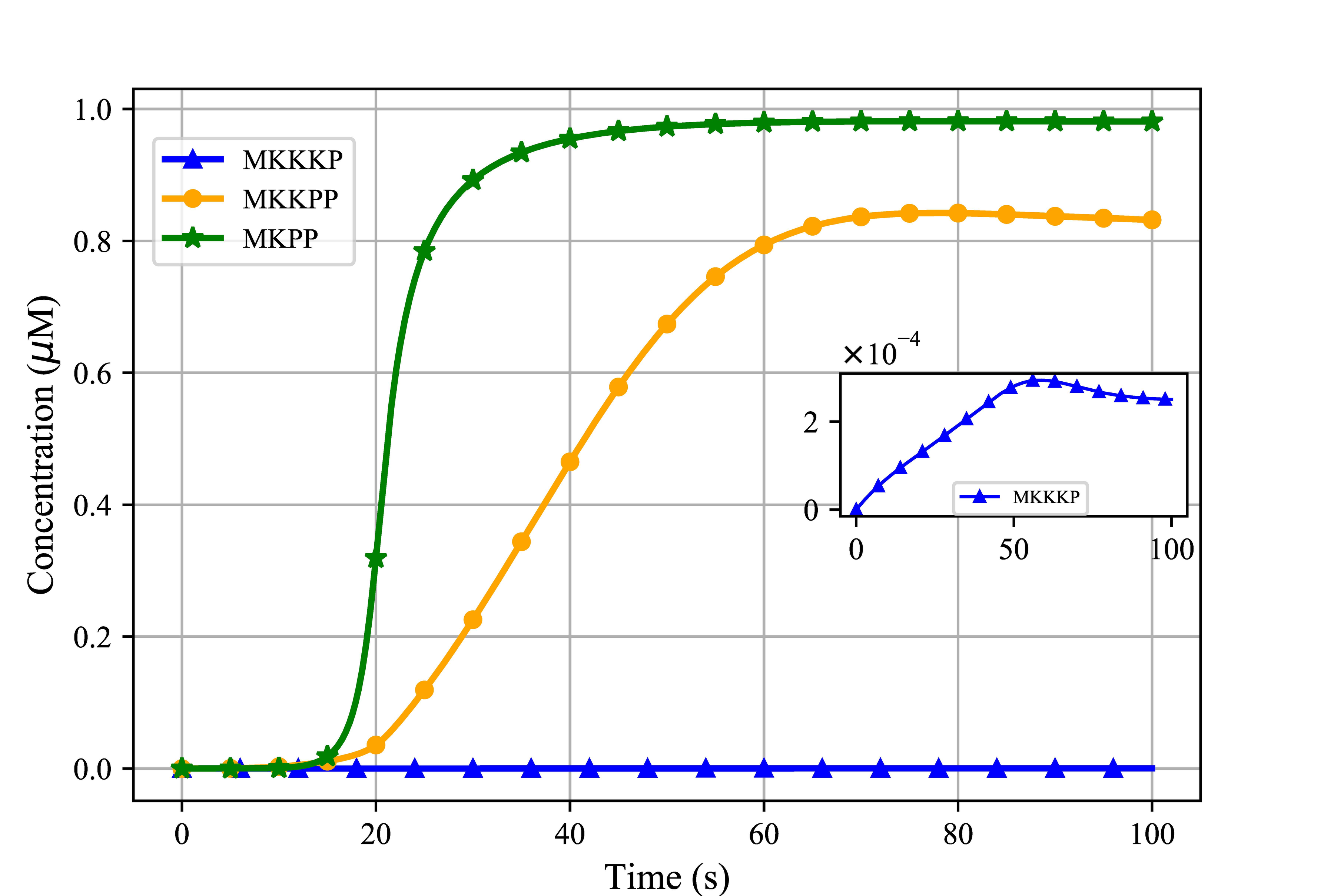

### S2 Fig

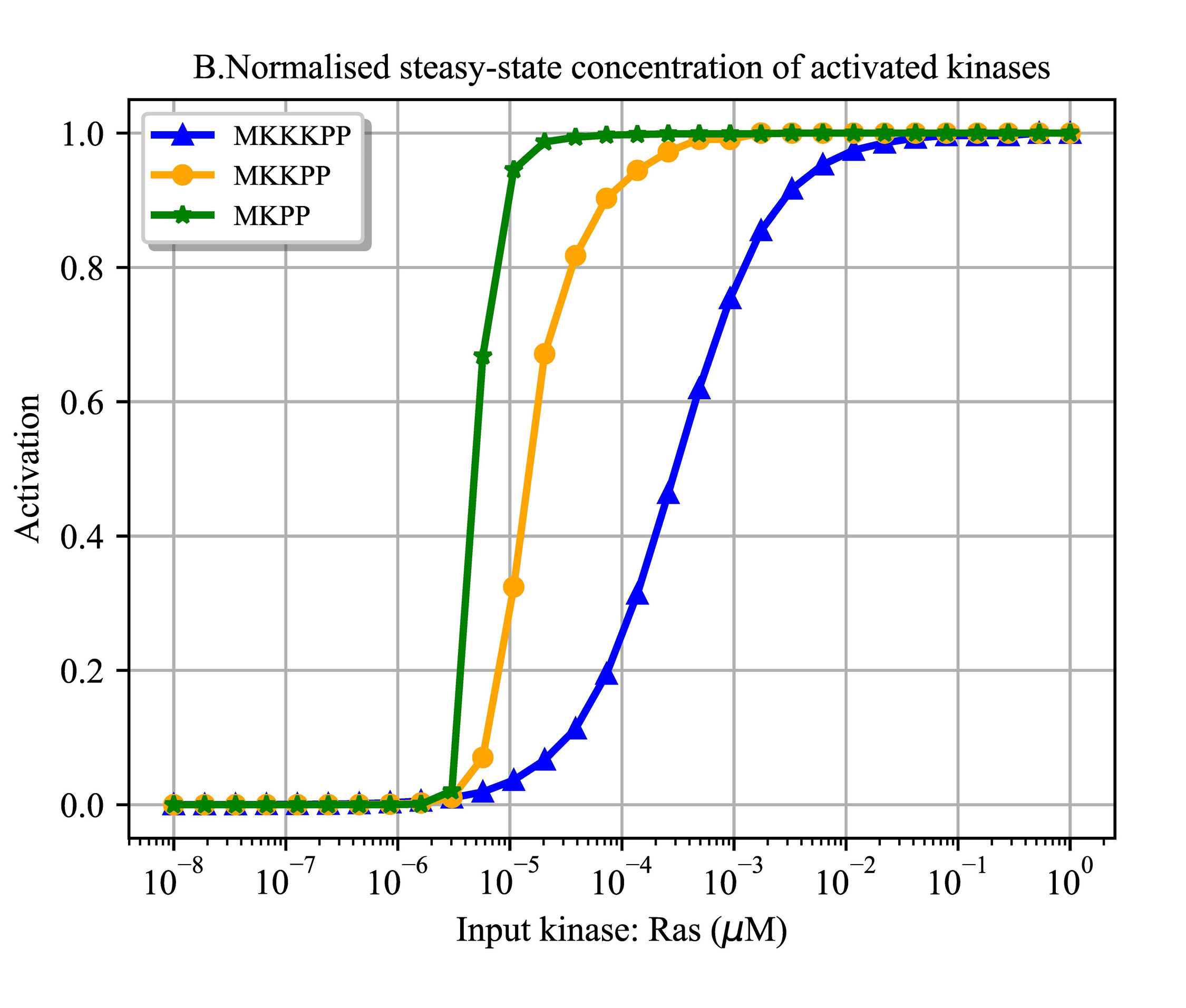
